# Systematic deletion of symmetrical *CFTR* exons reveals new therapeutic targets for exon skipping antisense oligonucleotides

**DOI:** 10.1101/2024.08.28.607949

**Authors:** Cecilia Pena-Rasgado, Elvia Manriquez, Miroslav Dundr, Robert J. Bridges, Michelle L. Hastings, Wren E. Michaels

## Abstract

There is a major need for therapeutics that treat diseases caused by pathogenic gene variants that disrupt protein open-reading frames. Splice-switching antisense oligonucleotides (ASOs) offer a potential solution by inducing the skipping of exons containing these variants, removing them from the mRNA and correcting the open-reading frame. Cystic fibrosis (CF), caused by disruption of the CF transmembrane regulator (*CFTR*) gene, is one such disease that has many chain-terminating variants, which are untreatable with standard protein-targeted modulator therapies. Using *CFTR* as a model, we demonstrate the utility of ASOs in engineering protein isoforms through exon skipping to rescue protein function disrupted by truncating variants. We functionally screened all CFTR isoforms generated by the deletion of symmetrical exons, which can be skipped without disrupting the open-reading frame. We identified exons that can be removed and produce CFTR isoforms that remain functionally responsive to modulators. We screened for ASOs that induce skipping of these exons and show that they recover CFTR function in airway cells derived from individuals with terminating *CFTR* variants. This study demonstrates that systematic functional analysis of in-frame exon-deleted protein isoforms can identify targets for ASO-based splice-switching therapies, a concept that can be broadly applied to any multi-exon protein-coding gene.

## Introduction

Premature termination codons (PTCs) generated by frameshift and nonsense mutations result in the synthesis of a truncated protein and trigger nonsense-mediated mRNA decay (NMD), largely abolishing functional protein production (1). Due to these multifaceted complications, treatments for diseases caused by terminating variants have been elusive. One therapeutic strategy for PTC-causing disease that has had clinical success uses splice-switching antisense oligonucleotides (ASOs) to induce exon skipping to eliminate pathogenic variants and/or reframe the mRNA transcript (2). This approach requires the exon targeted for ASO-induced skipping to be a symmetrical exon, defined as an exon in the same reading frame register as its flanking exons. Thus, when the symmetrical exon is spliced out, or skipped, the open reading frame is preserved. Symmetrical exon skipping is a common natural form of alternative splicing that can generate protein isoforms with altered functions (3). In the case of disease-associated variants in these exons, this mechanism can be harnessed using ASOs to remove PTCs and thereby stabilize the mRNA and recover protein expression. This approach can be therapeutic if the protein that is produced from the ASO-induced spliced transcript is at least partially functional. This strategy is the basis of the FDA-approved splice-switching antisense oligonucleotides targeting PTCs in Duchenne muscular dystrophy (DMD) (2).

To establish targets for exon-skipping strategies, an understanding of the function of protein isoforms encoded by mRNAs that lack a specific exon is required. In principle, rapid functional target identification can be achieved by systematic screening of all possible symmetrical exon deletions within a gene. Additionally, as this approach increases protein expression, small molecule drugs that target the protein can be screened for further rescue of protein function, as a potential combination therapy. Here, we demonstrate the utility of this screening approach for investigating treatment strategies for cystic fibrosis (CF), a genetic disorder affecting more than 100,000 people worldwide.

CF is an autosomal recessive disease caused by disruption of the cystic fibrosis transmembrane conductance regulator (*CFTR*) gene. *CFTR* encodes a transmembrane anion channel, the loss of which results in thick mucus buildup in multiple organs, most severely affecting the lungs and pancreas (4). There are thousands of different CF-associated *CFTR* variants that cause varying degrees of disease severity, based on their effect on expression and resulting protein function (5). For some variants, there are effective protein-targeted therapeutics approved to treat people with CF (pwCF). These include potentiator drugs, like Ivacaftor (VX-770), which enhance channel gating, and corrector drugs, used in combination with potentiators, which restore processing and trafficking of CFTR to the cell surface (6–8). The most prevalent treatment, Trikafta, is a combination of VX-770 and the correctors VX-661 and VX-445 (elexacaftor, tezacafter, ivacaftor: ETI) (7). These small molecule therapeutics, collectively referred to as highly effective modulator therapeutics (HEMTs), are effective in treating CF caused by more common variants, such as *CFTR-F508del*. However, because these drugs target the CFTR protein, they are not effective in treating CF caused by pathogenic variants that preclude protein production, such as those that produce PTCs. These types of variants account for ∼11% of all CF-causing variants and ∼62% of variants that are not eligible for treatment with current HEMTs (9–11).

For one such variant, *CFTR-W1282X*, we and others have reported on an effective ASO approach that corrects CFTR function by inducing the skipping of exon 23, a symmetrical exon which houses the variant (12–14). *CFTR* contains many other symmetrical exons that could be eliminated to remove variant-induced PTCs and NMD targets (**Figure 1A**). If the protein isoform lacking an exon retains some function and/or is amenable to rescue by HEMTs, ASO-mediated skipping of these exons has the potential to treat any truncating variants that have been identified within these *CFTR* exons (**Figure 1B**). Here, utilizing a systematic engineered protein screening approach, we identify CFTR symmetrical exons encoding amino acids in the C-terminus of the protein that can be removed and still produce a CFTR isoform with HEMT-responsive function. We show that ASO-induced skipping of these exons can induce these isoforms, providing the beginnings of a therapeutic path forward for treating CF caused by variants in these exons. Overall, we estimate ∼10% of CFTR variants that do not currently have effective treatment options may be responsive to this ASO-mediated exon skipping approach. These findings demonstrate the effectiveness of engineered protein screening to rapidly test for functional protein isoforms to guide strategic ASO design and assessment for exon skipping and functional protein production. This strategy offers a way to treat pathogenic terminating variants that can be broadly applied to any genetic disease.

**Figure 1.**
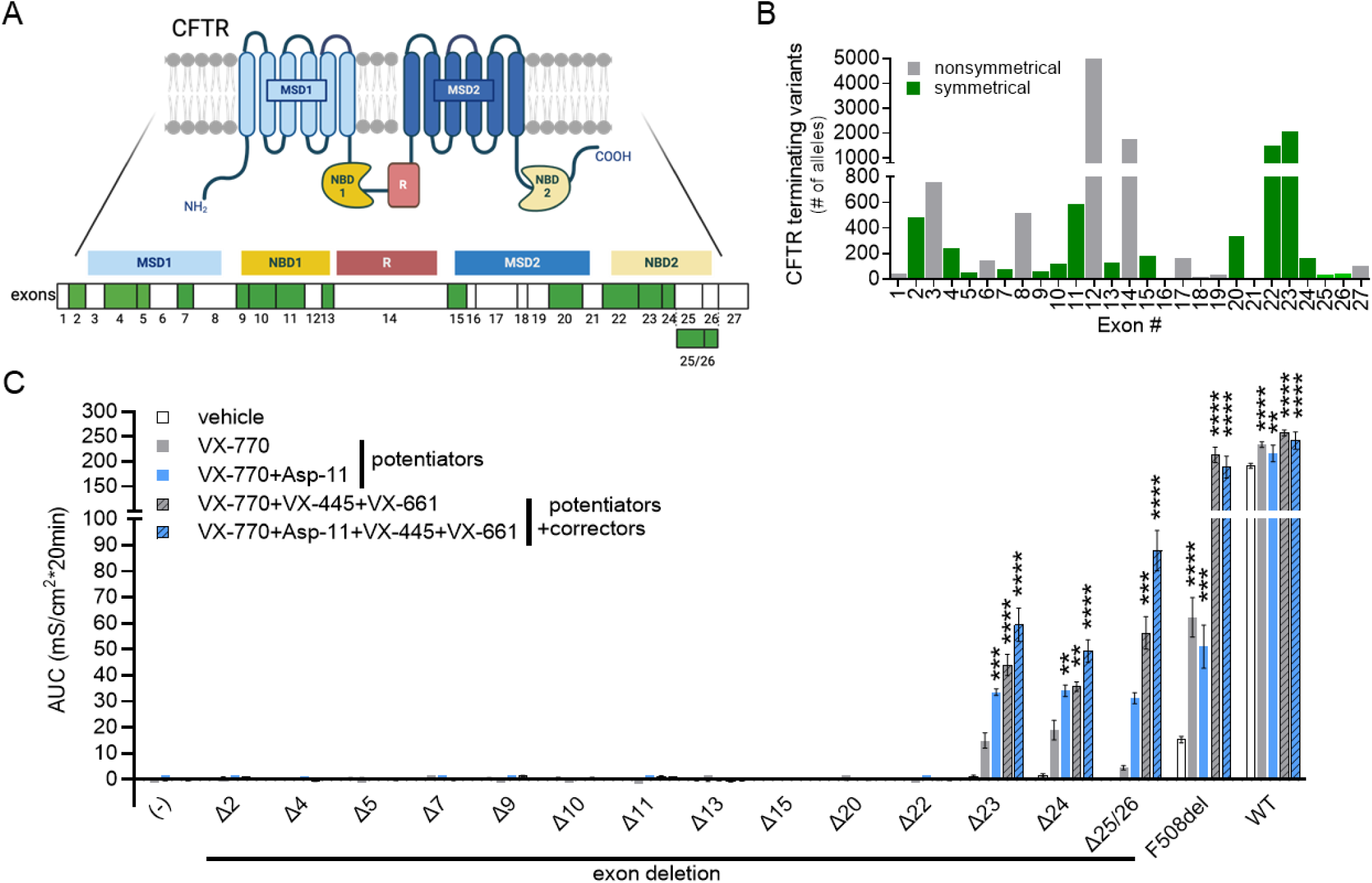
Systematic functional analysis of CFTR isoforms lacking symmetrical exons. (**A**) Schematic of CFTR protein in the cell membrane with protein domains aligned to CFTR exons. Symmetric exons are depicted in green. (**B**) Quantification of terminating variants found in each CFTR exon from pwCF registered in the CFTR2 database (data acquired April 7, 2023). Green bars represent variants in symmetric exons. (**C**) Measurement of chloride conductance in cells expressing CFTR or isoforms lacking individual symmetrical exons. Average area under the curve (AUC) was quantified for the forskolin or forskolin + potentiator(s) test periods from conductance traces (**Figure S1C**) measured in FRT cells, which lack endogenous CFTR, transfected with empty vector (-), CFTR-F508del, CFTR-WT, or each CFTR exon deletion construct. Cells were pre-treated with vehicle (DMSO), or VX-445+VX-661 (hashed bars) for 24 hours. Potentiators were added after forskolin addition during analysis (VX-770, gray bars, or VX-770+Asp-11, blue bars). Error bars are ±SEM; replicates are DMSO n=4, VX-770 n=3, VX-770+Asp-11 n=1, VX-770+VX-661+VX-445 n=3, VX-770+Asp-11+VX-661+VX-445 n=1 for all except: empty vector n=13,13,7,13,7; Δ5 n=5,3,2,2,2; Δ13 n=7,6,4,6,4; Δ22 n=3,4,3,4,3; Δ23 n=8,9,8,9,8; Δ24 n=5,6,5,6,5; Δ25/26 n=3(all); ΔF508 n=9,7,7,7,7; WT n=16,16,7,16,7 respectively; two-way ANOVA with Dunnett’s multiple comparison test compared to DMSO treatment within groups; **P<0.01, ***P<0.001, and ****P<0.0001.

## Materials and Methods

### Generation of exon deletion expression plasmids

CFTR-Δexon deletion plasmids were constructed from the synthetic *CFTR* high codon adaption index (HCAI) into the pcDNA3.1/Neo(+) vector using the Q5 Site-Directed Mutagenesis Kit (NEB) (**Table S1**) (15). All plasmid deletions were sequenced. Plasmids were reverse transfected into Fischer Rat Thyroid (FRT) cells with lipofectamine LTX (Thermo Fisher) and OptiMEM (Thermo Fisher). Clonal cell lines were selected with G418 (300 μg/ml) and maintained in media supplemented with G418 (150μg/ml).

### Cells and culture conditions

FRT cell lines were cultured in F12 Coon’s modification media (Sigma, F6636) supplemented with 10% fetal bovine serum (FBS) and 1% Penicillin-Streptomycin (PenStrep). T84 cells were cultured in Dulbecco’s Modified Eagle Medium/Nutrient Mixture F12 (DMEM/F12) media supplemented with 5% FBS and 1% PenStrep. Primary human bronchial epithelial (hBE) cells were acquired from the Compound Screening and Drug Discovery Core at RFUMS or the Cystic Fibrosis Foundation and thawed at passage (P) 1 or 2 for analysis. For functional studies cells were plated on 24-transwell filter plates (0.4 µM pore size, Polyester, Corning, catalog #CLS3397). FRT cells were grown in a liquid/liquid interface for 1 week. Primary cells were differentiated in an air/liquid interface for 5 weeks.

### Antisense oligonucleotide treatment

Antisense oligonucleotides were 25-mer phosphorodiamidate morpholino oligomers (PMO) formulated in sterile water (**Table S2**). A non-targeting PMO was used as a negative control, ASO-C (Gene Tools LLC).

T84 cells were transfected with ASOs in DMEM/F12 media supplemented with 5% FBS and 1% PenStrep using Endo-Porter (Gene-Tools, 6μl/ml) for 48 hrs. Primary human bronchial epithelial (hBE) cells were transfected after differentiation on filter plates as previously described (12) using a 1-hour hypo-osmotic shock. The cells were maintained in Dulbecco’s phosphate-buffered saline (DPBS) with ASO for 4 days until functional analysis.

### RNA isolation, RT-PCR, and real-time qPCR

RNA was extracted from cells using TRIzol (Thermo Fisher Scientific). Reverse transcription was performed on total RNA using the GoScript Reverse Transcription System with an oligo-dT primer (Promega). Splicing was analyzed by radiolabeled PCR of resulting cDNA using GoTaq Green (Promega) supplemented with α-^32^P-deoxycytidine triphosphate (dCTP). Primers for amplification are reported in **Table S3**. Reaction products were run on a 6% non-denaturing polyacrylamide gels and quantified using a Typhoon FLA 7000 phosphorimager (GE Healthcare) and ImageJ software.

Real-time qPCR was performed with PrimeTime Gene Expression Master Mix and PrimeTime qPCR probe assay kits for *CFTR* transcripts normalized to *HPRT1* (IDT, **Table S3**). All reactions were analyzed in triplicate. Real-time PCR was performed on an Applied Biosystems (ABI) ViiA 7 Real-Time PCR System and analyzed by the ΔΔCT method.

### Protein isolation and immunoblot analysis

Cell lysates were prepared using NP-40 lysis buffer (1% Igepal, 150mM NaCl, 50mM Tris-HCl pH7.6) supplemented with 1× protease inhibitor cocktail (Sigma-Aldrich, cat #11836170001). Protein concentration was measured using a Coomassie (Bradford) protein assay (Thermo Fisher, cat #23200). Cell lysates were prepared with 4X NuPage LDS sample buffer (Invitrogen) for a final concentration of 1-2 mg/ml, incubated at 37°C for 5 minutes, and loaded into a 4-12% Bis-Tris gel (Criterion) for SDS-PAGE separation in 1X MOPS buffer. Samples were transferred in 1X CAPS buffer to polyvinylidene difluoride membranes. Protein was detected using primary anti-CFTR antibodies 570 or 596 (Riordan lab UNC, Cystic Fibrosis Foundation, diluted 1:1000) and anti-β-actin (C4, Santa Cruz Biotechnology, diluted 1:2000). Secondary anti-mouse poly-HRP was added for chemiluminescent detection with HRP substrate (CFTR:Femto, β-actin:Classico; Immobulin, Sigma). Signal was quantified with ImageJ software.

### Automated conductance/equivalent current assay

FRT cells lines and primary hBE cells were treated with VX-445+VX-661 (1μM + 3μM for hBEs or 3 μM + 3.5 μM for FRTs, Selleckchem) or vehicle (0.1% or 0.2% DMSO) at 37°C for 24 hours. Cells were switched from culture media to HEPES-buffered (pH 7.4) F12 Coon’s modification running media (Sigma, F6636) apically and basolaterally and allowed to equilibrate for one hour at 37°C without CO_2_. For functional measurements the transepithelial voltage (Vt) and resistance (Rt) measurements were recorded at 37°C with a 24-channel TECC robotic system (EP Design, Belgium) as previously described (12,16). Benzamil (6 μM), forskolin (10 μM), VX-770 (1 μM) with or without ASP-11 (20 μM), and Inh-172 (20 μM) or bumetanide (20 μM) additions were added sequentially. Conductance (Gt) was calculated by the reciprocal of the recorded Rt (Gt=1/Rt) and plotted as conductance traces (**Figure S1**). Equivalent currents from hBE donors were assessed similarly and calculated using Ohm’s law (Ieq=Vt/Rt). Area under the curve measurements of forskolin, forskolin + VX-770, or forskolin + VX-770 + ASP-11, were calculated using a one-third trapezoidal rule for each test period using Microsoft Excel.

### Immunofluorescence Imaging

FRT cell lines grown on high precision glass 12 mm circle coverslips (No. 1.5H) (Paul Marlenfeld GmbH) were washed with PBS and fixed with 4% paraformaldehyde in PBS for 10 min at room temperature. Cells were permeabilized with 0.2% Triton X-100 in PBS for 5 min on ice. Subsequently, the cells were incubated in 3% bovine serum albumin for 20 min. Cells were incubated with the monoclonal anti-CFTR antibody (570, Riordan lab UNC, 1:500) and rabbit polyclonal anti-Claudin antibody (1:100, Zymed) for one hour. Cells were then incubated with goat anti-mouse secondary antibody conjugated with Alexa 488 and goat anti–rabbit secondary antibody conjugated with Alexa 594 (Invitrogen, 1:1000) for 1 hour and mounted using ProLong Gold antifade mounting reagent with DAPI (Life Technologies). Cells were observed on Zeiss Elyra structured illumination microscope (SIM) using a 63 × 1.40 NA objective using z-90 nm sectioning. In the axial direction, channel alignment was performed using 100 nm tetraspeck beads. Final images were prepared using Zeiss Zen software.

### Statistics

Statistical analyses were performed using GraphPad PRISM 10.2.3 or Microsoft Excel. Specific statistical tests used in each experiment are reported in the figure legends.

## Results

### Systematic deletion of *CFTR* symmetrical exons reveals functional protein isoforms

*CFTR* has 13 symmetrical exons that can be removed without disrupting the protein open reading frame. Additionally, the reading frame is maintained when exons 25 and 26 are eliminated together (**Figure 1A**). We and others have previously shown that ASO-induced skipping of exon 23 results in a partially functional CFTR protein when co-treated with established HEMTs (12–14). To identify additional pathogenic variants that could potentially be treated using this therapeutic approach, we analyzed CFTR protein lacking each of the other symmetrical exons. We created a series of *CFTR* plasmids with individual deletions of each of the 13 exons and both exons 25 and 26 together (CFTR-Δex) and analyzed *CFTR* expression and function after stable transfection into Fischer Rat Thyroid (FRT) cells, which do not express endogenous *CFTR*. CFTR-Δex isoform expression was confirmed by RT-PCR (**Figure S2A)**.

FRT CFTR-Δex cell lines were grown to confluence on trans-well filter plates to assess the protein isoforms’ cAMP-activated conductance capabilities across the plasma membrane via forskolin stimulation. We also screened for responsiveness to HEMTs including ETI and the novel potentiator Asp-11, which has been shown to work in synergy with VX-770 to further increase the function of CFTR protein isoforms, especially those that disrupt the second nucleic binding domain (NBD2) of CFTR (17,18). None of the expressed CFTR isoforms had measurable conductance in the FRT cells in the absence of HEMTs (white bars, **Figure 1C, S1**). Treatment with VX-770 alone (grey bars, **Figure 1C**) did not result in measurable conductance, however, co-potentiation with Asp-11 in combination with VX-770 (blue bars, **Figure 1C**) resulted in significant forskolin-induced conductance from CFTR-Δ23 and CFTR-Δ24 compared to vehicle-treated cells. Cells expressing CFTR-Δ23, CFTR-Δ24, and CFTR-Δ25/26 had significant conductance following ETI treatment (grey hashed bars, **Figure 1C**) and all three deletion isoforms responded most robustly when treated with a combination of Asp-11 and ETI (blue hashed bars, **Figure 1C**).

Conductance activity did not correlate with expression as immunoblot analysis of the C-terminal deletion protein isoforms did not show significant expression differences above the other deletion isoforms (**Figure S2B**). Preliminary analysis of protein processing showed an increase in mature CFTR Band C protein after ETI treatment in comparison to the immature Band B, indicative of recovery of proper CFTR folding and trafficking. These results provide evidence that this activity was specific to modulator effects on the isoforms (**Figure S2B**). Additionally, immunofluorescence analysis of cells expressing CFTRΔex23, 24 and 25/26 showed appropriate localization of the expressed proteins to the cell membrane after ETI treatment (**Figure S3**). Taken together, these findings indicate that CFTR isoforms lacking exons 23, 24, or 25 and 26 together can transport chloride across the membrane when treated with HEMTs, making them good candidates for combined HEMT and ASO-mediated exon skipping therapies.

### Splice-switching ASOs induce skipping of CFTR symmetrical exons

We next tested splice-switching ASOs that target each symmetrical exon for skipping. A series of ASOs for each exon were designed to base-pair with the canonical acceptor and donor splice sites or putative exonic splicing enhancer sequences and further selected for their predicted binding affinity (**Figure 2A**). ASOs were tested for their ability to induce exon skipping of endogenous *CFTR-WT* pre-mRNA by transfection into T84 cells. RT-PCR analysis of RNA isolated after 48 hrs of incubation identified at least one ASO that was effective at skipping each target exon, except exon 26 where combination with exon 25 skipping ASOs was necessary (**Figure 2B, S4A**).

**Figure 2.**
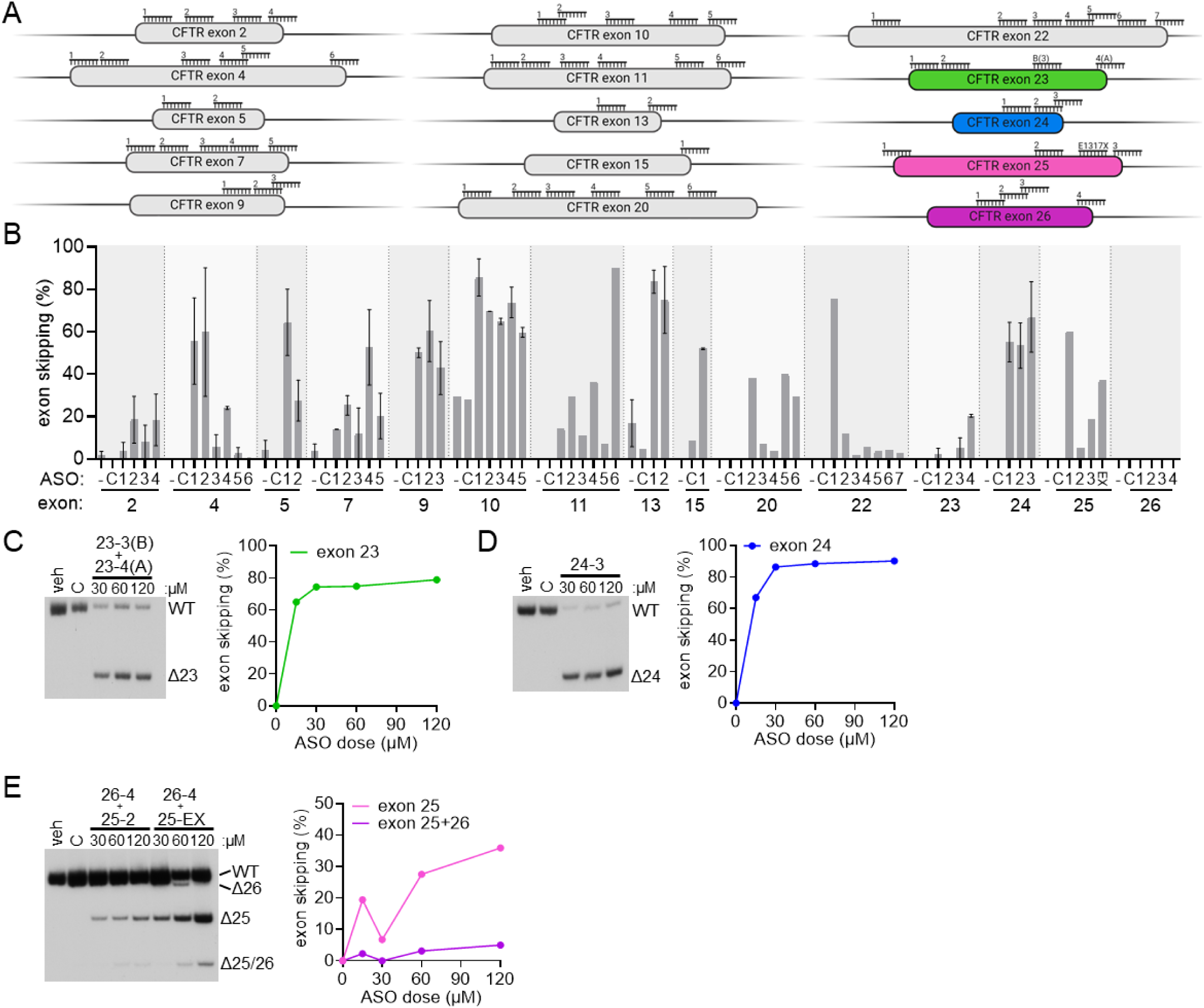
Antisense oligonucleotides induce skipping of symmetrical exons. (**A**) Schematic of ASOs aligned to each *CFTR* exon targeted for skipping. (**B**) RT-PCR quantification of exon skipping [Δ/(full-length+cryptic+Δ)]*100 for each ASO (15µM). (**C-E**) (left) RT-PCR analysis of RNA isolated from T84 cells transfected with vehicle (veh), control ASO (C, 120µM), or ASO-23A+ASO-23B (**C**) ASO-24-3 (**D**) or ASO-26-4 + ASO-25-2 or ASO-25-EX (**E**) at indicated doses. (right) The percent of transcripts with exon skipping was plotted in relation to ASO dose. Primer pairs within exons flanking each exon targeted for skipping were used for analysis (**Table S3**).

Our functional screen indicated that only isoforms lacking exon 23, 24 or 25/26 are responsive to HEMT and thus, we focused on ASOs targeting these exons. An ASO that base-pairs to the 5’ splice site region of exon 23 (ASO-23A) induces skipping of exon 23, but also causes splicing to a cryptic splice site that results in an out-of-frame CFTR mRNA. Additional treatment with an ASO that base-pairs at the cryptic splice site (ASO-23B) eliminates cryptic splicing and greatly enhances skipping of exon 23 in a dose-dependent manner (**Figure 2C**) (12). Likewise, a robust dose-dependent increase in exon 24 was observed following treatment with a single ASO, ASO-24-3 (**Figure 2D**). Treatment with ASO-25-2 or 25-EX, which targets the *CFTR-E1371X* variant, alone or in combination with ASO-26-4 (ASO-25EX+26) resulted in the skipping of exon 25 or 26 individually or both exons 25 and 26 together (Δ25/26) (**Figure 2E, S4B**). Though dual skipping of exon 25 and 26 was not induced at a high level, higher dosing of ASO-25EX+26 provided some improvement (**Figure 2E**). Because elimination of exon 25 alone would lead to PTCs in the penultimate exon 26, which may escape NMD (1), ASO treatment could provide sufficient splice modulation to increase CFTR mRNA and protein for further rescue with HEMTs.

### Functional CFTR isoforms generated by ASO-induced symmetrical exon skipping in cells derived from CF patient airways

We recently reported that treatment with ASOs that induce exon 23 skipping can rescue CFTR function in cells homozygous for *CFTR-W1282X* (12). This rescue required treatment with ETI. Here, we tested whether compound heterozygous cells expressing only a single copy of *CFTR-W1282X* along with another variant that is associated with minimally functional CFTR could produce sufficient CFTR-Δex23 to provide a functional benefit in genotypes that aren’t currently approved for HEMTs. We also tested co-potentiation with Asp-11, to assess whether greater functional rescue could be achieved by modulator optimization. CFTR function was assessed via equivalent current analysis in human bronchial epithelial cells (hBE) isolated from pwCF.

Cells from donors expressing two copies of *CFTR-W1282X* or with one copy and another nonsense variant, *CFTR-R553X* or *CFTR-R1162X,* were treated with vehicle, ASO-C, or ASO-23AB with or without Asp-11 and ETI. In all donors, treatment with ASO-23AB resulted in a significant increase in chloride currents over the control ASO when treated with ETI, and co-treatment with Asp-11 resulted in an even greater ASO-induced rescue of CFTR function (**Figure 3A**). In donor cells with *CFTR-W1282X* and *CFTR-F508del*, ASO-23AB treatment had little effect on the rescue already achieved by ETI alone, similar to previous results (12). Interestingly in these cells, ASO-23AB and co-potentiator treatment with VX-770 and Asp-11 together resulted in significant rescue of CFTR function, but Asp-11 addition with ETI lowered functional recovery (**Figure S5A**). This effect on ETI-dependent function was also observed in cells homozygous for *CFTR-F508del*, where ASO-23AB treatment had no effect on modulator rescue (**Figure S5A**). Co-potentiation alone provided a significant rescue in cells from the CFTR-W1282X/CFTR-R553X donor (**Figure 3A**). These results suggest that HEMT optimization could both provide significant rescue for minimal function variants and further improve the ASO-induced CFTR-Δex23 isoform function. The results also caution that effects on the second CFTR allele should be considered, as differential modulator effects may occur.

**Figure 3.**
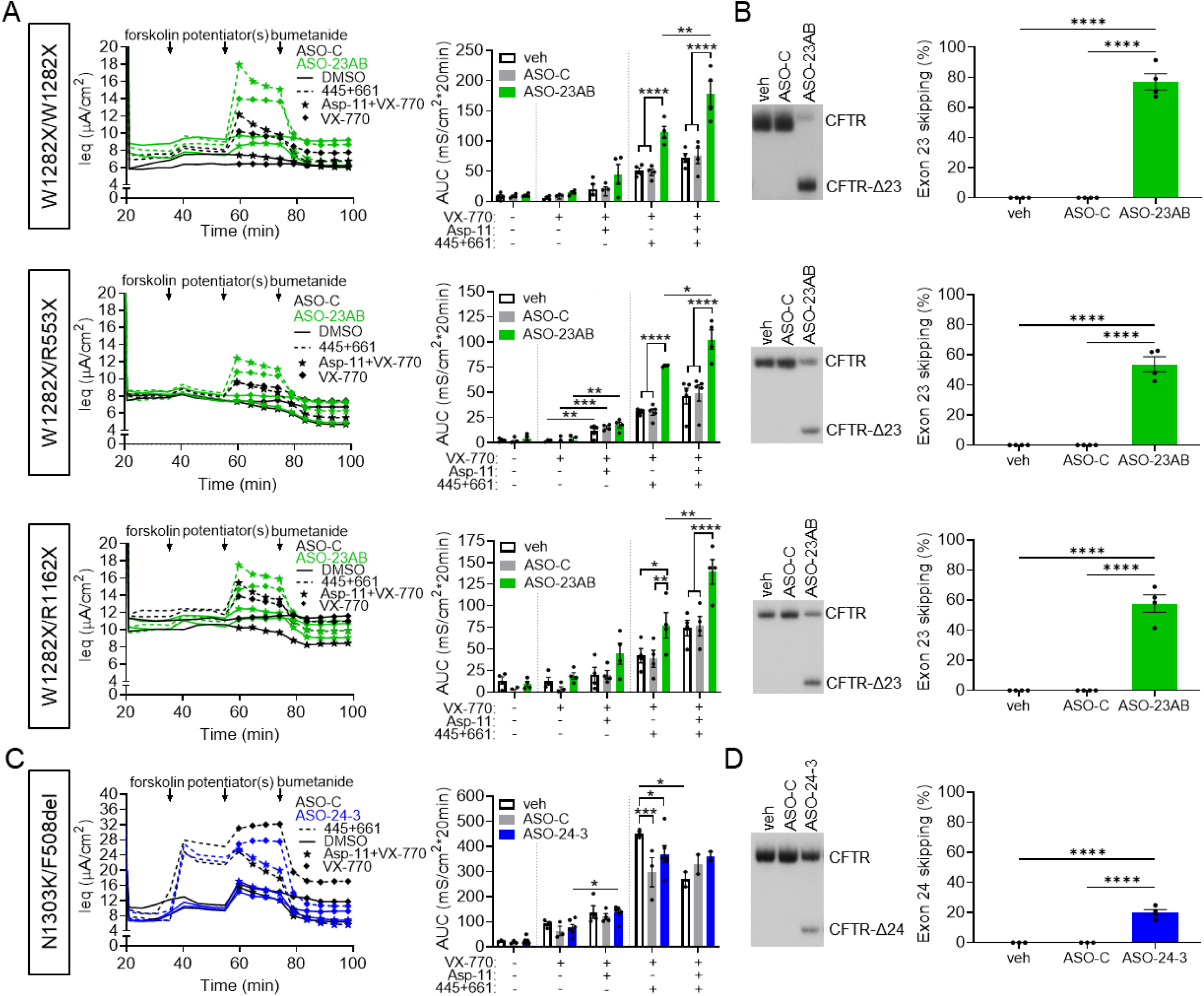
ASO treatment induces exon skipping and rescues CFTR function in primary human bronchial epithelial (hBE) cells isolated from pwCF when used with HEMTs. (**A**) (left) Equivalent current (Ieq) traces of primary hBE cells from CF donors expressing *CFTR-W1282X*. The genotype of each donor is indicated on the left. Cells were transfected with ASO-C (80 μM, black), or ASO-23AB (40 μM each, green) and treated with vehicle (DMSO, solid lines) or VX-445+VX-661 (dashed lines) for 24 hours before functional analysis. Forskolin, potentiator [VX-770 (diamond) or VX-770+Asp-11(star)], and bumetanide additions are indicated. (right) Average AUC of the current traces was quantified for the forskolin or forskolin+potentiator(s) test periods for each treatment group. Error bars are ±SEM; two-way ANOVA with Tukey’s multiple comparison test within groups to assess ASO effects or Šídák’s multiple comparison test between groups to assess ASP-11 effects; *P<0.05, **P<0.01, ***P<0.001, and ****P<0.0001. (**B**) (left) RT-PCR analysis of exon 23 splicing in hBE cells indicated to the left. (right) Quantification of the percent of mRNA with exon 23 skipping for each treated donor. Error bars are ±SEM; one-way ANOVA with Tukey’s multiple comparisons test; ****P<0.0001. (**C**) (left) Equivalent current (Ieq) traces of primary hBE cells from a CF donor compound heterozygous for CFTR-N1303K and CFTR-F508del. Cells were transfected with ASO-C (40-160 μM, black), or ASO-24-3 (40-160 μM, blue) and treated as in **A**. (right) Average AUC of the current trace was quantified for the forskolin or forskolin+potentiator(s) test periods for each treatment group. Error bars are ±SEM; two-way ANOVA with Tukey’s multiple comparisons test within groups to assess ASO effects or Šídák’s multiple comparison test between groups to assess ASP-11 effects; *P<0.05 and ***P<0.001. (**D**) (left) RT-PCR analysis of exon 24 splicing in hBE cells. (right) Quantification of exon 24 skipping. Error bars are ±SEM; one-way ANOVA with Tukey’s multiple comparisons test; ****P<0.0001.

We have reported that ASO-induced exon skipping could have partial allele specificity for *CFTR-W1282X* due to the elimination of splicing factor binding sites (12). Similarly here, ASO treatment induced ∼80% exon 23 skipping in cells homozygous for *CFTR-W1282X* (**Figure 3B**). In compound heterozygous cells, ASO treatment induced ∼30-60% exon skipping (**Figure 3B**, **S5B**). In cells homozygous for *CFTR-F508del*, ASO-23AB treatment induced less than 20% skipping of the wildtype exon 23 (**Figure S5B**), providing strong evidence that exon 23 containing the W1282X sequence is more susceptible to ASO-23AB-induced skipping than the wildtype sequence. These results suggest that ASO-induced exon 23 skipping could be therapeutic for pwCF expressing at least one copy of *CFTR-W1282X* and not eligible for, or responsive to, current therapeutics.

Skipping of exon 24 also results in a CFTR protein that is functional when treated with HEMTs (**Figure 1**). Thus, we evaluated the effect of ASO-induced exon 24 skipping on CFTR function in patient donor cells. Unfortunately, airway cells from a donor with a terminating variant in exon 24 were not available. Instead, we tested ASO-24-3 in hBE cells derived from a person with the *CFTR-N1303K* variant, a CF-associated missense variant in exon 24 that results in a protein with defective biogenesis and gating abnormalities. This variant has shown some response to HEMTs but is not currently FDA-approved for HEMT treatment (17,19,20). Given the activity of the CFTR-Δ24 isoform (**Figure 1**), we hypothesized that CFTR-Δ24 would have greater activity than *CFTR-N1303K* and, because of this enhanced activity, ASO-induced skipping of exon 24 would result in elevated currents in the donor cells. Compound heterozygote hBE cells expressing *CFTR-N1303K* and *CFTR-F508del* were transfected with ASO-24-3 and treated with ETI and Asp-11. Compared to the control ASO, there was little effect of ASO-24-3 treatment on ETI rescue of CFTR activity and Asp-11 had no additional effect (**Figure 3C**). This result suggests that ASO-induced skipping of exon 24 will not provide a therapeutic benefit over ETI to individuals with *CFTR-N1303K* and *CFTR-F508del*. However, ASO-24-3 and co-potentiator treatment did result in a significant rescue over VX-770 alone (**Figure 3C**) despite inducing only modest (∼20%) exon 24 skipping (**Figure 3D**), suggesting that further optimization of ASO-induced skipping could result in greater activity, particularly in the context of other CF variants that are not responsive to ETI. Nonetheless, *CFTR-N1303K* is an imperfect model for exon skipping therapies as there is no evidence of mRNA degradation. Further testing in donor cells expressing variants in exon 24 that destabilize CFTR mRNA is needed to assess the efficacy of ASO-induced exon 24 skipping.

### A potential functional CFTR isoform generated by ASO-induced multiexon skipping

Similar to CFTR-Δ23 and CFTR-Δ24, CFTR-Δ25/26 retains conductance when treated with ETI and is responsive to co-potentiation with Asp-11 (**Figure 1C**). ASO-25-EX alone induces robust exon 25 skipping (**Figure 2B**), which could be therapeutic if the downstream PTC in the penultimate exon does not activate NMD. The combination of ASO-26-4 with ASO-25EX (ASO-25EX+26) results in modest skipping of both exons 25 and 26 (**Figure 2E**). To test these ASOs further, hBE cells that are compound heterozygous for *CFTR-E1371X*, a PTC in exon 25, and *CFTR-F508del*, were treated with ASO-25-EX alone or ASO-25EX+26 and assessed for CFTR activity in the presence or absence of ETI and Asp-11. ASO treatment had little effect on activity without ETI treatment (**Figure 4A**). With ETI, ASO-25EX+26 showed no additional rescue and ASO-25EX resulted in slightly lower function compared to the control ASO. As seen before in cells expressing *CFTR-F508del*, Asp-11 treatment reduced ETI response but did show a significant increase in function in combination with ASO-25EX+26 treatment compared to control treated cells, which may suggest that Asp-11 has some specificity for CFTR-Δ25/26, which could lead to elevated function in genotypes heterozygous with variants other than *CFTR-F508del* (**Figure 4A**). ASO-25EX treatment resulted in ∼10-20% exon 25 skipping alone or in combination with ASO-26-4 and ASO-25EX+26 induced a low level of skipping of both exons 25 and 26 (**Figure 4B**). Allele-specific qPCR analysis revealed a trend of increased *CFTR-E1371X* mRNA expression with ASO-25-EX+26 treatment despite the minimal skipping, suggesting that ASO treatment may effectively stabilize mRNA produced from *CFTR-E1371X*, presumably due to the removal of the PTC in the ASO-induced CFTR25/26 isoform (**Figure 4C**). The upward trend in mRNA levels, and the modest effect on CFTR function with ETI and Asp-11 treatment suggests that further optimization of ASO-induced exon 25 and 26 skipping could be beneficial to people with terminating variants in exons 25 or 26.

**Figure 4.**
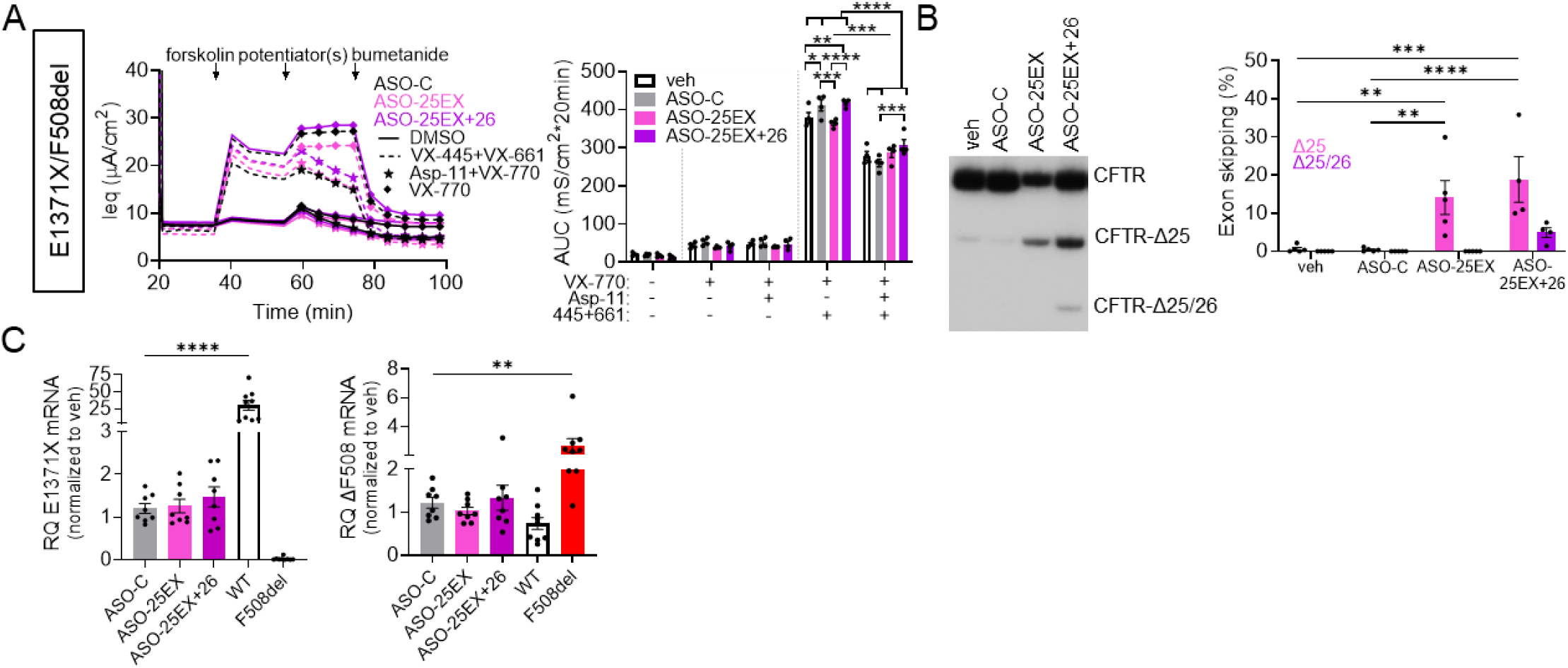
An ASO cocktail induces multi-exon skipping of CFTR exons 25 and 26 in primary human bronchial epithelial (hBE) cells isolated from pwCF.(**A**) (left) Equivalent current (Ieq) traces of primary hBE cells from a CF donor compound heterozygous for CFTR-F508del and CFTR-E1371X. Cells were transfected with ASO-C (40-240 µM, black), ASO-25 (40-120 µM, pink), or ASO-25+ASO-26 (40-120 µM each, purple) and treated with vehicle (DMSO, solid lines) or VX-445+VX-661 (dashed lines) for 24 hours. Forskolin, potentiator [VX-770 (diamond) or VX-770+ASP-11 (star)], and bumetanide additions are indicated. (right) Average AUC of the current traces was quantified for the forskolin or forskolin+potentiator(s) test periods for each treatment group. Error bars are ±SEM two-way ANOVA with Tukey’s multiple comparison test within groups to assess ASO effects or Šídák’s multiple comparison test between groups to assess ASP-11 effects; *P<0.05, **P<0.01, ***P<0.001, and ****P<0.0001. (**B**) (left) RT-PCR analysis of exon 25 or exon 25/26 splicing in hBE cells from A (right) Quantification of exon 25 or 25/26 skipping. Error bars are ±SEM; two-way ANOVA with Tukey’s multiple comparisons test **P<0.01, ***P<0.001, and ****P<0.0001. (**C**) RT-qPCR analyses of total (left) *CFTR-E1371X* mRNA (exon 11-12) or (right) *CFTR-F508del* mRNA (exon 11-12) from treated hBE cells compared to RNA from a non-CF or homozygous CFTR-F508del donor. Error bars are ±SEM; one-way ANOVA with Dunnet’s multiple comparisons test to ASO-C; **P<0.01 and ****P<0.0001.

### Inactive CFTR produced from ASO-induced exon 22 skipping confirms functional limits of exon-deleted isoforms

The systematic screening of protein isoforms engineered by ASO-induced exon skipping revealed that CFTR C-terminal isoforms lacking amino acids encoded by exons 23, 24, or 25 and 26, but not exon 22 or any other upstream exon, are functional as chloride channels when treated with ETI (**Figure 1**). To confirm the results found in this functional screen, we tested ASO-induced skipping of exon 22 using ASO-22-1 in hBE cells derived from two pwCF expressing *CFTR-R1162X (*compound heterozygous with *W1282X)* or *CFTR-c.3658delC* (p.K1177SfsX15) (compound heterozygous with *F508del*). In the *CFTR-c.3658delC* cells, ASO-22-1 treatment did not improve activity relative to that already achieved by ETI treatment alone and Asp-11 had no additional effects (**Figure 5A**). In the *CFTR-R1162X* cells, ETI and Asp-11 treatment significantly improved CFTR function in control treated cells, likely due to the effect on *CFTR-W1282X*, but treatment with ASO-22-1 eliminated this response (**Figure 5A**). Lack of functional recovery was not due to insufficient skipping of exon 22 as treatment with ASO-22-1 resulted in ∼30% exon 22 skipping in *CFTR-3659delC* cells and ∼70% in *CFTR-R1162X* cells (**Figure 5B**). These results, along with the conductance activity measured from CFTR-Δ22 in FRT cells, suggest that ASO-induced skipping of exon 22 does not result in an active CFTR isoform and that the protein domains encoded by exons 1-22, are likely essential for functional activity and rescue with HEMTs. These results define the boundaries of any potential exon-skipping therapy for CF as sequences downstream of exon 22 and suggest that truncation or deletions after CFTR exon 22 could provide a therapeutic benefit for all variants effecting expression in CFTR exons 23-27.

**Figure 5.**
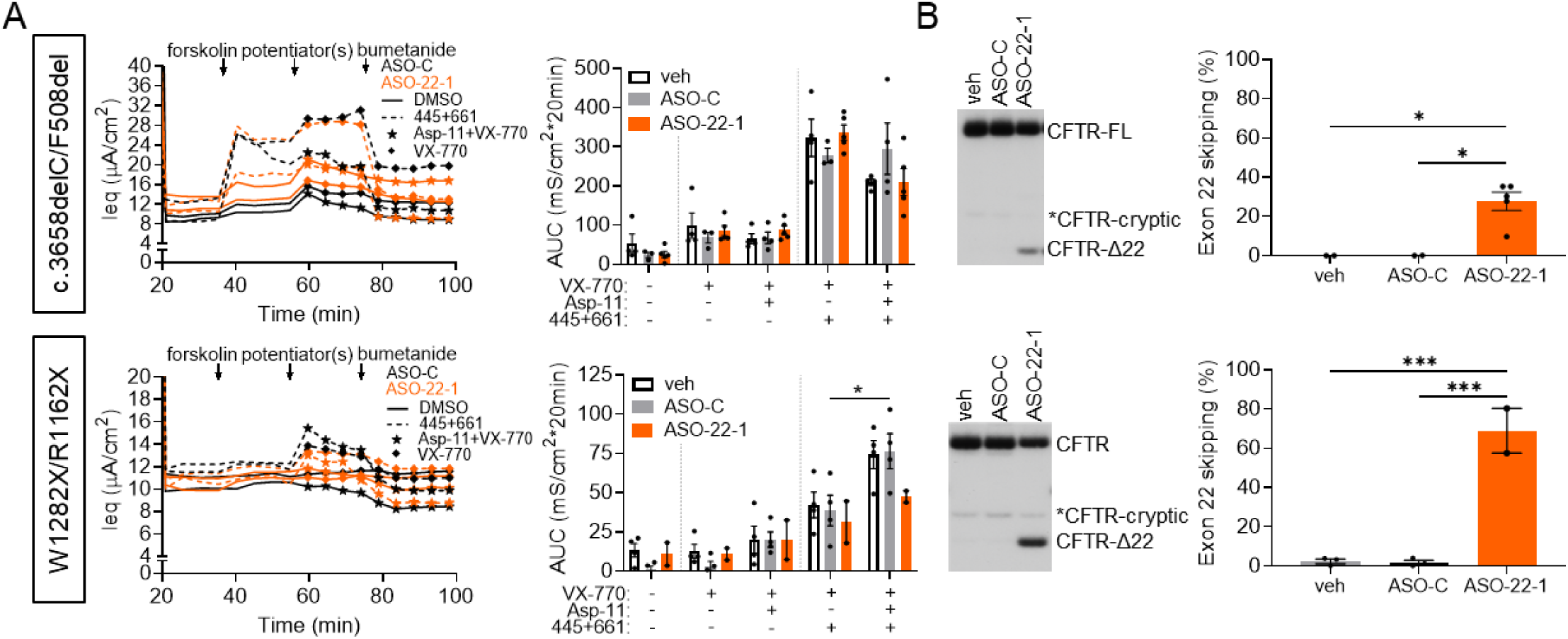
ASO treatment induces exon 22 skipping but does not rescue CFTR function in primary human bronchial epithelial (hBE) cells isolated from a CF individual expressing terminating variants. (**A**) (left) Equivalent current (Ieq) traces of primary hBE cells from CF donors expressing at least one nonsense variant in exon 22. The genotype of each donor is indicated at the left. Cells were transfected with ASO-C (40-160 µM, black), or ASO-22-1 (40-160 µM, orange) and treated with vehicle (DMSO, solid lines) or VX-445+VX-661 (dashed lines) for 24 hours. Forskolin, potentiator [VX-770 (diamond) or VX-770+Asp-11 (star)], and bumetanide additions are indicated. (right) Average AUC of the current traces was quantified for the forskolin or forskolin+potentiator(s) test periods for each treatment group. Error bars are ±SEM; within groups to assess ASO effects or Šídák’s multiple comparison test between groups to assess ASP-11 effects; *P<0.05. (**B**) (left) RT-PCR analysis of exon 22 splicing from cells in **A**. (right) Quantification of exon 22 skipping. Error bars are ±SEM; one-way ANOVA with Tukey’s multiple comparisons test; *P<0.05 and ***P<0.001.

## Discussion

Identifying targets of ASO-mediated exon skipping that can restore functional potential to an inactivated protein is relevant to therapeutic development for any disease caused by pathogenic variants that result in PTCs and destabilize the mRNA and/or truncate the protein, including CF. To this end, our study demonstrates the systematic testing of each so-called skippable (i.e. symmetrical) exon of the CFTR gene for its functional dispensability and identifies four exons at the 3’ end of the transcript as viable targets for restoring function by ASO-induced exon skipping.

We and others have previously demonstrated the potential of ASO-induced exon skipping for treating CF caused by *CFTR-W1282X* (12–14). Additional testing of this approach in patient cells expressing genotypes not currently eligible for HEMTs provides strong evidence that combining ASO treatment with HEMTs will provide therapeutic benefit to these patients and others that have terminating variants in exon 23 (**Figure 3**). Our testing of additional exon deletion isoforms in comparison to the CFTR-Δex23 protein product, demonstrates that CFTR-Δex24 and CFTR-Δex25/26 have comparable levels of function and HEMT response (**Figure 1**), providing evidence that this strategy could be beneficial to terminating variants in additional C-terminal exons. Our subsequent screening identifies ASOs that induce skipping of these exons (**Figure 2**).

Further ASO design and testing in CF patient cells is needed to advance these approaches. However, screening is limited by a lack of patient cells with required genotypes needed for functional testing, limiting our analysis to cells with less-than-ideal CFTR genotypes for definitively evaluating therapeutic benefit. As such, no additional benefit over ETI was observed with ASO-mediated exon 24 skipping in patient cells expressing *CFTR-N1303K* with *CFTR-F508del* (**Figure 3**). However, because *CFTR-N1303K* has not been shown to compromise CFTR expression, it is a sub-optimal target for this strategy. Furthermore, the tested donor cells co-expression with *CFTR-F508del* makes it difficult to assess ASO+HEMT efficacy against the *CFTR-F508del* ETI response. Significant ASO-specific rescue was seen with co-potentiation treatment alone, suggesting modulated CFTR-Δ24 has some benefit. However, further testing in patients with terminating variants in exon 24 and other undruggable genotypes is needed. Regarding exon 25/26 skipping, responses from patient cells expressing *CFTR-E1371X* revealed a significant response from ASO-25EX+26 compared to ASO-C treated cells with Asp-11 addition to ETI. However, again, ASO-specific functional rescue of *CFTR-E1371X* directly was difficult to assess with *CFTR-F508del*. These results, as well as a trend of increased expression of *CFTR-E1371X* RNA despite low levels of exon 25/26 skipping, suggest that ASO treatment could be promising for terminating variants in these exons with further optimization (**Figure 4**).

In addition to ASOs, a number of alternative therapeutic strategies are being explored to improve stability and translation of mRNA with terminating mutations, though currently none have been successfully implemented in patients. Current strategies involve approaches to block NMD and/or increase translational readthrough of PTCs to recover full-length protein expression (21–27). One downfall of such candidate therapeutics is that, global dysregulation of these processes could be detrimental if specificity for the PTC-containing transcript is not achieved. Gene-specific, ASO-mediated suppression of NMD offers a more tailored approach that may overcome these issues but are limited because of the potential need for more than one ASO in order to target multiple sites on the mRNA that trigger NMD (28,29). Editing and CFTR replacement strategies are other approaches that hold promise providing that delivery and control of off-target editing can be achieved (30–35). The pursuit of all technologies for treating disease caused by PTCs is undoubtedly the way to develop the best and most efficacious treatments. ASOs offer some advantages for design and development that present the potential for a relatively short development timeline, which is especially critical to individuals with these variants that currently have no good treatment options, including a large population of pwCF.

ASOs may offer some advantage over other strategies as a validated therapeutic platform, proven largely safe in humans. In-frame exon skipping strategies utilizing ASOs have been successfully applied to address PTCs in DMD (2) and explored in other disease models (36–42). ASOs also offer a potential advantage over larger therapeutic cargo in delivery to target tissues. ASO delivery to the respiratory system, one of the primary targets for CF therapeutics, has been demonstrated clinically for asthma and other inflammatory lung conditions (43) and most recently as an antiviral approach for Sars-CoV-2 (44,45). Naked ASOs have been successfully delivered to the lung and enter multiple cell types including CFTR expressing epithelial cells (46–49), though questions about whether the targeted cell-types are relevant for CF treatment remain (50). ASOs have long-lasting effects in primary hBE cells isolated from pwCF (16), and could have similar durability *in vivo* considering ASO activity for other indications can last weeks to months in terminally differentiated cells (51,52). Lung epithelia may turnover relatively slowly, with estimates of 6-18 months (53).

Aerosolization of ASOs as a delivery modality has been demonstrated and targeted *CFTR* transcripts in both a CF-like lung disease model in mice and monkeys as well as pwCF (48,49,54,55). However, ASOs for CF have not yet advanced beyond clinical trials. Additionally, people with terminating variants tend to have severe disease phenotypes in other organs, so systemic delivery of ASOs must also be considered (56). Intravenous and subcutaneous ASO injections are currently used to treat other diseases, demonstrating the potential for systemic delivery of ASOs for CF (57–59). Successful ASO delivery to the affected organs in pwCF and target engagement with *CFTR* will be essential to therapeutic success.

In addition to therapeutic discovery, exploring the effects of exon deletions can provide information on protein function that can inform and facilitate future drug discovery and development. Our systematic screening of CFTR exon deletion isoforms indicates that the C-terminal exons are not essential for the fundamental function of CFTR when HEMTs are applied, a result confirmed by ASO testing in CF airway cells. Previous studies have shown that proteins with C-terminal truncations up to amino acid 1217 retain partial surface expression and function and are responsive to HEMTs (60–63). The domains encoded by exons 23, 24, and 25/26 encompass parts of NBD2 (NBD2=1203-1440, Δ23=Δ1240-1291, Δ24=Δ1292-1321, and Δ2526=Δ1321-1413), and the retained function of these exon deletions, but not exons upstream, is consistent with these reports (**Figure 1C**). CFTR-Δ23 and CFTR-Δ24 also retain critical sites in NBD2 for ATP-binding and NBD dimerization (64), which are eliminated in CFTR-Δ25/26. Similar conductance levels between the isoforms (**Figure 1C**) suggest that, if this region is critical for dimerization, NBD dimerization is not critical for proper CFTR function with HEMT treatment. Furthermore, removal of PTCs in these exons allows for translation to the C-terminus, which includes amino acids important for stability, gating, and localization (63,65–67). Overall, our functional data shows that elimination of exon 23, 24, or 25/26 results in CFTR protein responsive to HEMT rescue and suggests that ASO-mediated exon skipping of CFTR after exon 22 could be a therapeutic pathway. In addition, these results provide evidence that pwCF with splicing mutations that eliminate exons 23-26 are good candidates for ETI treatment if not currently approved for its use.

Co-potentiation with Asp-11, previously shown to increase the activity of other variants that are unresponsive to ETI, further enhanced CFTR isoform function (**Figure 1,3-5**) (17,18). This result suggests that new modulator discovery focused on enhancing function from CFTR deletion isoforms could provide additional benefit to ASO-induced rescue of expression, especially considering that the binding sites of the modulators are eliminated by some deletions. For example, the deletion of exon 20 eliminates a potential VX-770 binding pocket and, though not responsive to VX-770, might be responsive to alternative potentiators (69). Class I correctors bind in the first transmembrane domain (TMD1) and could be affected by several of the deletions, including exon 9 which has been previously explored for exon skipping and showed no response to the class 1 corrector VX-809 (70,71). The differing binding sites of VX-445 and VX-661 improve the efficacy of ETI and similar approaches to modulator design may be beneficial for each deletion construct (72).

Taken together, our results not only inform CF drug development, but also have important implications for treating other diseases caused by terminating variants. Treating such variants by eliminating the exons they reside in and increasing productive protein expression could be a broadly applied therapeutic strategy. Our investigation into the effects of symmetrical exon removal on CFTR expression and function provides a roadmap for the systematic evaluation of symmetrical exon deletion, revealing additional ASO therapeutic targets for CF drug development and instructing on a therapeutic strategy for discovering treatment approaches for other diseases.

## Acknowledgments

We thank the CFF for providing the patient hBE cells and CFTR antibodies. We also thank Marc Anderson for providing Asp-11. We also thank the M.L.H., R.J.B., and W.E.M. laboratories for assistance with experiments and insightful discussions, in particular James Hogan and Dr. Jessica Centa for their help in editing the manuscript. This study was funded by grants from the CFF 006048G223 (W.E.M. and M.L.H.), 00022F220 (W.E.M) and HASTING19GO (M.L.H and R.J.B), and NIH OD010662 to the Midwest Proteome Center.

---

**Figure S1.** Representative current traces of CFTR constructs analyzed in Figure 1C.

**Figure S2.** Expression of CFTR exon deletion constructs. **(A)** RT-PCR analysis of *CFTR* RNA collected from FRT cells transfected with each CFTR exon deletion construct, compared to CFTR-WT. **(B)** (top) Immunoblot analysis of CFTR protein, bands C and B, isolated from FRT cells stably transfected with empty vector, CFTR-F508del, CFTR-WT, or each CFTR exon deletion construct. Cells were treated with DMSO or VX-445+VX-661. β-actin was used as a loading control. (middle) Total CFTR protein (band C+band B/ β-actin was quantified. (bottom) The ratio of CFTR (band C / band B) isoforms was quantified. Replicates are n=2 and experimental groups are connected by a line.

**Figure S3.** Cellular localization of CFTR and the exon skipped isoforms. Immunofluorescent images of FRT cells transfected with empty vector, CFTR-WT, CFTR-F508del, CFTR-Δ22, CFTR-Δ23, CFTR-Δ24, or CFTR-Δ25/26. Cells were incubated with a CFTR-specific monoclonal antibody (green) and a rabbit Claudin polyclonal antibody (magenta). Regions of interest at the cell membrane were enlarged to show co-localization of CFTR at the cell membrane (white).

**Figure S4.** Spliced-switching ASOs induce skipping of symmetrical CFTR exons. (**A**) RT-PCR analysis of T84 cells transfected with vehicle, control ASO (ASO-C), or CFTR exon targeting ASOs. Amplicons representing cryptically spliced CFTR is indicated by * and expected exon skipping for each ASO is labelled with Δ on the right side of the gel. Quantification of gels are shown in **Figure 2B**. ASOs that induced the highest level of skipping are highlighted in green. (**B**) RT-PCR analysis of T84 cells transfected with vehicle, control ASO (ASO-C), or CFTR exon 25 and 26 targeting ASOs. Exon skipping is quantified as [Δ/(full-length+Δ)]*100. Primers flanking the exon were used for analysis (**Table S3**).

**Figure S5.** ASO treatment induces *CFTR-W1282X* exon skipping but does not increase function above approved HEMTs when expressed with CFTR-F508del (**A**) (left) Equivalent current (Ieq) traces of primary hBE cells from CF donors. The genotype of each donor is indicated on the left. Cells were transfected with ASO-C (80 μM, black), or ASO-23AB (40 μM each, green) and treated with vehicle (DMSO, solid lines) or VX-445+VX-661 (dashed lines) for 24 hours. Forskolin, potentiator [VX-770 (diamond) or VX-770+Asp-11(star)], and bumetanide additions are indicated. (right) Average AUC of the current traces was quantified for the forskolin or forskolin+potentiator(s) test periods for each treatment group. Error bars are ±SEM two-way ANOVA with Tukey’s multiple comparison within groups to assess ASO effects or Šídák’s multiple comparison test between groups to assess ASP-11 effects; *P<0.05, **P<0.01. (**B**) (left) RT-PCR analysis of exon 23 splicing in hBE cells from A. (right) Quantification of percent exon 23 skipping for each treated donor. Error bars are ±SEM; one-way ANOVA with Tukey’s multiple comparisons test; *P<0.05, **P<0.01, ****P<0.0001.

**Table S1.** Primers flanking *CFTR* symmetrical exons used to generate the *CFTR* exon deletion constructs.

**Table S2.** Antisense oligonucleotide sequences used in this study.

**Table S3.** Primers and probes used for PCR and qPCR analysis.

## References

1. Das, R. and Panigrahi, G.K. (2024) Messenger RNA Surveillance: Current Understanding, Regulatory Mechanisms, and Future Implications. Mol Biotechnol.

2. Gupta, S., Sharma, S.N., Kundu, J., Pattanayak, S. and Sinha, S. (2023) Morpholino oligonucleotide-mediated exon skipping for DMD treatment: Past insights, present challenges and future perspectives. J Biosci, 48.

3. Magen, A. and Ast, G. (2005) The importance of being divisible by three in alternative splicing. Nucleic Acids Res, 33, 5574–5582.

4. Cheng, S.H., Gregory, R.J., Marshall, J., Paul, S., Souza, D.W., White, G.A., O’Riordan, C.R. and Smith, A.E. (1990) Defective intracellular transport and processing of CFTR is the molecular basis of most cystic fibrosis. Cell, 63, 827–834.

5. Guo, J., Garratt, A. and Hill, A. (2022) Worldwide rates of diagnosis and effective treatment for cystic fibrosis. J Cyst Fibros, 21, 456–462.

6. Van Goor, F., Hadida, S., Grootenhuis, P.D.J., Burton, B., Stack, J.H., Straley, K.S., Decker, C.J., Miller, M., McCartney, J., Olson, E.R., et al. (2011) Correction of the F508del-CFTR protein processing defect in vitro by the investigational drug VX-809. Proceedings of the National Academy of Sciences of the United States of America, 108, 18843–18848.

7. Ferreira, F.C., Buarque, C.D. and Lopes-Pacheco, M. (2024) Organic Synthesis and Current Understanding of the Mechanisms of CFTR Modulator Drugs Ivacaftor, Tezacaftor, and Elexacaftor. Molecules, 29.

8. Van Goor, F., Hadida, S., Grootenhuis, P.D.J., Burton, B., Cao, D., Neuberger, T., Turnbull, A., Singh, A., Joubran, J., Hazlewood, A., et al. (2009) Rescue of CF airway epithelial cell function in vitro by a CFTR potentiator, VX-770. Proceedings of the National Academy of Sciences, 106, 18825-18830.

9. Bear, C.E. (2020) A Therapy for Most with Cystic Fibrosis. Cell, 180, 211–211.

10. De Boeck, K., Zolin, A., Cuppens, H., Olesen, H.V. and Viviani, L. (2014) The relative frequency of CFTR mutation classes in European patients with cystic fibrosis. Journal of cystic fibrosis: official journal of the European Cystic Fibrosis Society, 13, 403–409.

11. Cutting, G.R., Castellani, C., Corey, M., Lewis, M.H., Penland, C.M., Raraigh, K.S., Rommens, J.M. and Sosnay, P.R. (2013), Cftr2.

12. Michaels, W.E., Pena-Rasgado, C., Kotaria, R., Bridges, R.J. and Hastings, M.L. (2022) Open reading frame correction using splice-switching antisense oligonucleotides for the treatment of cystic fibrosis. Proc Natl Acad Sci U S A, 119, e2114886119.

13. Kim, Y.J., Sivetz, N., Layne, J., Voss, D.M., Yang, L., Zhang, Q. and Krainer, A.R. (2022) Exon-skipping antisense oligonucleotides for cystic fibrosis therapy. Proc Natl Acad Sci U S A, 119.

14. Oren, Y.S., Avizur-Barchad, O., Ozeri-Galai, E., Elgrabli, R., Schirelman, M.R., Blinder, T., Stampfer, C.D., Ordan, M., Laselva, O., Cohen-Cymberknoh, M. et al. (2022) Antisense oligonucleotide splicing modulation as a novel Cystic Fibrosis therapeutic approach for the W1282X nonsense mutation. J Cyst Fibros, 21, 630–636.

15. Shah, K., Cheng, Y., Hahn, B., Bridges, R., Bradbury, N.A. and Mueller, D.M. (2015) Synonymous Codon Usage Affects the Expression of Wild Type and F508del CFTR. Journal of Molecular Biology, 427, 1464–1479.

16. Michaels, W.E., Bridges, R.J. and Hastings, M.L. (2020) Antisense oligonucleotide-mediated correction of CFTR splicing improves chloride secretion in cystic fibrosis patient-derived bronchial epithelial cells. Nucleic Acids Res, 48, 7454–7467.

17. Phuan, P.W., Tan, J.A., Rivera, A.A., Zlock, L., Nielson, D.W., Finkbeiner, W.E., Haggie, P.M. and Verkman, A.S. (2019) Nanomolar-potency ‘co-potentiator’ therapy for cystic fibrosis caused by a defined subset of minimal function CFTR mutants. Sci Rep, 9, 17640.

18. Phuan, P.W., Son, J.H., Tan, J.A., Li, C., Musante, I., Zlock, L., Nielson, D.W., Finkbeiner, W.E., Kurth, M.J., Galietta, L.J. et al. (2018) Combination potentiator (’co-potentiator’) therapy for CF caused by CFTR mutants, including N1303K, that are poorly responsive to single potentiators. J Cyst Fibros, 17, 595–606.

19. Sadras, I., Kerem, E., Livnat, G., Sarouk, I., Breuer, O., Reiter, J., Gileles-Hillel, A., Inbar, O., Cohen, M., Gamliel, A. et al. (2023) Clinical and functional efficacy of elexacaftor/tezacaftor/ivacaftor in people with cystic fibrosis carrying the N1303K mutation. J Cyst Fibros, 22, 1062–1069.

20. Destefano, S., Gees, M. and Hwang, T.-C. (2018) Physiological and pharmacological characterization of the N1303K mutant CFTR. Journal of Cystic Fibrosis, 17, 573–581.

21. Venturini, A., Borrelli, A., Musante, I., Scudieri, P., Capurro, V., Renda, M., Pedemonte, N. and Galietta, L.J.V. (2021) Comprehensive Analysis of Combinatorial Pharmacological Treatments to Correct Nonsense Mutations in the CFTR Gene. Int J Mol Sci, 22.

22. Ko, W., Porter, J.J., Sipple, M.T., Edwards, K.M. and Lueck, J.D. (2022) Efficient suppression of endogenous CFTR nonsense mutations using anticodon-engineered transfer RNAs. Mol Ther Nucleic Acids, 28, 685–701.

23. Albers, S., Allen, E.C., Bharti, N., Davyt, M., Joshi, D., Perez-Garcia, C.G., Santos, L., Mukthavaram, R., Delgado-Toscano, M.A., Molina, B. et al. (2023) Engineered tRNAs suppress nonsense mutations in cells and in vivo. Nature, 618, 842–848.

24. Carollo, P.S., Tutone, M., Culletta, G., Fiduccia, I., Corrao, F., Pibiri, I., Di Leonardo, A., Zizzo, M.G., Melfi, R., Pace, A., et al. (2023) Investigating the Inhibition of FTSJ1, a Tryptophan tRNA-Specific 2’-O-Methyltransferase by NV TRIDs, as a Mechanism of Readthrough in Nonsense Mutated CFTR. Int J Mol Sci, 24.

25. Leubitz, A., Vanhoutte, F., Hu, M.-Y., Porter, K., Gordon, E., Tencer, K., Campbell, K., Banks, K. and Haverty, T. (2021) A Randomized, Double-Blind, Placebo-Controlled, Multiple Dose Escalation Study to Evaluate the Safety and Pharmacokinetics of ELX-02 in Healthy Subjects. Clinical pharmacology in drug development, 10, 859-869.

26. Temaj, G., Telkoparan-Akillilar, P., Nuhii, N., Chichiarelli, S., Saha, S. and Saso, L. (2023) Recoding of Nonsense Mutation as a Pharmacological Strategy. Biomedicines, 11.

27. Keenan, M.M., Huang, L., Jordan, N.J., Wong, E., Cheng, Y., Valley, H.C., Mahiou, J., Liang, F., Bihler, H., Mense, M. et al. (2019) Nonsense-mediated RNA Decay Pathway Inhibition Restores Expression and Function of W1282X CFTR. Am J Respir Cell Mol Biol, 61, 290–300.

28. Kim, Y.J., Nomakuchi, T., Papaleonidopoulou, F., Yang, L., Zhang, Q. and Krainer, A.R. (2022) Gene-specific nonsense-mediated mRNA decay targeting for cystic fibrosis therapy. Nat Commun, 13, 2978.

29. Nomakuchi, T.T., Rigo, F., Aznarez, I. and Krainer, A.R. (2016) Antisense oligonucleotide-directed inhibition of nonsense-mediated mRNA decay. Nature Biotechnology, 34, 164–166.

30. Lomunova, M.A. and Gershovich, P.M. (2023) Gene Therapy for Cystic Fibrosis: Recent Advances and Future Prospects. Acta Naturae, 15, 20–31.

31. Li, C., Liu, Z., Anderson, J., Liu, Z., Tang, L., Li, Y., Peng, N., Chen, J., Liu, X., Fu, L. et al. (2023) Prime editing-mediated correction of the CFTR W1282X mutation in iPSCs and derived airway epithelial cells. PLoS One, 18, e0295009.

32. Mention, K., Cavusoglu-Doran, K., Joynt, A.T., Santos, L., Sanz, D., Eastman, A.C., Merlo, C., Langfelder-Schwind, E., Scallan, M.F., Farinha, C.M. et al. (2023) Use of adenine base editing and homology-independent targeted integration strategies to correct the cystic fibrosis causing variant, W1282X. Hum Mol Genet, 32, 3237-3248.

33. Belbellaa, B., Reutenauer, L., Messaddeq, N., Monassier, L. and Puccio, H. (2020) High Levels of Frataxin Overexpression Lead to Mitochondrial and Cardiac Toxicity in Mouse Models. Mol Ther Methods Clin Dev, 19, 120–138.

34. Rowe, S.M., Zuckerman, J.B., Dorgan, D., Lascano, J., McCoy, K., Jain, M., Schechter, M.S., Lommatzsch, S., Indihar, V., Lechtzin, N. et al. (2023) Inhaled mRNA therapy for treatment of cystic fibrosis: Interim results of a randomized, double-blind, placebo-controlled phase 1/2 clinical study. J Cyst Fibros, 22, 656–664.

35. Vaidyanathan, S., Kerschner, J.L., Paranjapye, A., Sinha, V., Lin, B., Bedrosian, T.A., Thrasher, A.J., Turchiano, G., Harris, A. and Porteus, M.H. (2024) Investigating adverse genomic and regulatory changes caused by replacement of the full-length CFTR cDNA using Cas9 and AAV. Mol Ther Nucleic Acids, 35, 102134.

36. Molinari, E., Ramsbottom, S.A., Srivastava, S., Booth, P., Alkanderi, S., McLafferty, S.M., Devlin, L.A., White, K., Gunay-Aygun, M., Miles, C.G. et al. (2019) Targeted exon skipping rescues ciliary protein composition defects in Joubert syndrome patient fibroblasts. Sci Rep, 9, 10828.

37. Leier, A., Moore, M., Liu, H., Daniel, M., Hyde, A.M., Messiaen, L., Korf, B.R., Selvakumaran, J., Ciszewski, L., Lambert, L. et al. (2022) Targeted exon skipping of NF1 exon 17 as a therapeutic for neurofibromatosis type I. Mol Ther Nucleic Acids, 28, 261–278.

38. Ablinger, M., Lettner, T., Friedl, N., Potocki, H., Palmetzhofer, T., Koller, U., Illmer, J., Liemberger, B., Hainzl, S., Klausegger, A. et al. (2021) Personalized Development of Antisense Oligonucleotides for Exon Skipping Restores Type XVII Collagen Expression in Junctional Epidermolysis Bullosa. International Journal of Molecular Sciences, 22, 3326.

39. Lee, J.J.A., Maruyama, R., Duddy, W., Sakurai, H. and Yokota, T. (2018) Identification of Novel Antisense-Mediated Exon Skipping Targets in DYSF for Therapeutic Treatment of Dysferlinopathy. Molecular Therapy - Nucleic Acids, 13, 596–604.

40. Yamamura, T., Horinouchi, T., Adachi, T., Terakawa, M., Takaoka, Y., Omachi, K., Takasato, M., Takaishi, K., Shoji, T., Onishi, Y. et al. (2020) Development of an exon skipping therapy for X-linked Alport syndrome with truncating variants in COL4A5. Nature Communications, 11.

41. Centa, J.L., Stratton, M.P., Pratt, M.A., Osterlund Oltmanns, J.R., Wallace, D.G., Miller, S.A., Weimer, J.M. and Hastings, M.L. (2023) Protracted CLN3 Batten disease in mice that genetically model an exon-skipping therapeutic approach. Mol Ther Nucleic Acids, 33, 15–27.

42. Rodriguez-Polo, I. and Behr, R. (2022) Exploring the Potential of Symmetric Exon Deletion to Treat Non-Ischemic Dilated Cardiomyopathy by Removing Frameshift Mutations in TTN. Genes (Basel), 13.

43. Liao, W., Dong, J., Peh, H.Y., Tan, L.H., Lim, K.S., Li, L. and Wong, W.F. (2017) Oligonucleotide Therapy for Obstructive and Restrictive Respiratory Diseases. Molecules, 22.

44. Zhu, C., Lee, J.Y., Woo, J.Z., Xu, L., Nguyenla, X., Yamashiro, L.H., Ji, F., Biering, S.B., Van Dis, E., Gonzalez, F., et al. (2022) An intranasal ASO therapeutic targeting SARS-CoV-2. Nat Commun, 13, 4503.

45. Yu, A.M. and Tu, M.J. (2022) Deliver the promise: RNAs as a new class of molecular entities for therapy and vaccination. Pharmacol Ther, 230, 107967.

46. Brinks, V., Lipinska, K., de Jager, M., Beumer, W., Button, B., Livraghi-Butrico, A., Henig, N. and Matthee, B. (2019) The Cystic Fibrosis-Like Airway Surface Layer Is not a Significant Barrier for Delivery of Eluforsen to Airway Epithelial Cells. J Aerosol Med Pulm Drug Deliv, 32, 303–316.

47. Crosby, J.R., Zhao, C., Jiang, C., Bai, D., Katz, M., Greenlee, S., Kawabe, H., McCaleb, M., Rotin, D., Guo, S. et al. (2017) Inhaled ENaC antisense oligonucleotide ameliorates cystic fibrosis-like lung disease in mice. J Cyst Fibros, 16, 671–680.

48. Zhao, C., Crosby, J., Lv, T., Bai, D., Monia, B.P. and Guo, S. (2019) Antisense oligonucleotide targeting of mRNAs encoding ENaC subunits alpha, beta, and gamma improves cystic fibrosis-like disease in mice. J Cyst Fibros, 18, 334–341.

49. Ozeri-Galai, E., Friedman, L., Barchad-Avitzur, O., Markovetz, M.R., Boone, W., Rouillard, K.R., Stampfer, C.D., Oren, Y.S., Hill, D.B., Kerem, B. et al. (2023) Delivery Characterization of SPL84 Inhaled Antisense Oligonucleotide Drug for 3849 + 10 kb C-> T Cystic Fibrosis Patients. Nucleic Acid Ther, 33, 306–318.

50. Shin, M., Chan, I.L., Cao, Y., Gruntman, A.M., Lee, J., Sousa, J., Rodriguez, T.C., Echeverria, D., Devi, G., Debacker, A.J. et al. (2022) Intratracheally administered LNA gapmer antisense oligonucleotides induce robust gene silencing in mouse lung fibroblasts. Nucleic Acids Res, 50, 8418–8430.

51. Luu, K.T., Norris, D.A., Gunawan, R., Henry, S., Geary, R. and Wang, Y. (2017) Population Pharmacokinetics of Nusinersen in the Cerebral Spinal Fluid and Plasma of Pediatric Patients With Spinal Muscular Atrophy Following Intrathecal Administrations. J Clin Pharmacol, 57, 1031–1041.

52. Migliorati, J.M., Liu, S., Liu, A., Gogate, A., Nair, S., Bahal, R., Rasmussen, T.P., Manautou, J.E. and Zhong, X.B. (2022) Absorption, Distribution, Metabolism, and Excretion of US Food and Drug Administration-Approved Antisense Oligonucleotide Drugs. Drug Metab Dispos, 50, 888–897.

53. Rawlins, E.L. and Hogan, B.L. (2008) Ciliated epithelial cell lifespan in the mouse trachea and lung. Am J Physiol Lung Cell Mol Physiol, 295, L231–234.

54. Drevinek, P., Pressler, T., Cipolli, M., De Boeck, K., Schwarz, C., Bouisset, F., Boff, M., Henig, N., Paquette-Lamontagne, N., Montgomery, S., et al. (2020) Antisense oligonucleotide eluforsen is safe and improves respiratory symptoms in F508DEL cystic fibrosis. J Cyst Fibros, 19, 99–107.

55. Fajac, I. and De Boeck, K. (2017) New horizons for cystic fibrosis treatment. Pharmacol Ther, 170, 205–211.

56. Orenti, A., Pranke, I., Faucon, C., Varilh, J., Hatton, A., Golec, A., Dehillotte, C., Durieu, I., Reix, P., Burgel, P.R. et al. (2023) Nonsense mutations accelerate lung disease and decrease survival of cystic fibrosis children. J Cyst Fibros, 22, 1070–1079.

57. Ardizzone, S., Bevivino, G. and Monteleone, G. (2016) Mongersen, an oral Smad7 antisense oligonucleotide, in patients with active Crohn’s disease. Therap Adv Gastroenterol, 9, 527–532.

58. Quemener, A.M., Centomo, M.L., Sax, S.L. and Panella, R. (2022) Small Drugs, Huge Impact: The Extraordinary Impact of Antisense Oligonucleotides in Research and Drug Development. Molecules, 27.

59. Sheng, L., Rigo, F., Bennett, C.F., Krainer, A.R. and Hua, Y. (2020) Comparison of the efficacy of MOE and PMO modifications of systemic antisense oligonucleotides in a severe SMA mouse model. Nucleic Acids Res, 48, 2853–2865.

60. Cui, L., Aleksandrov, L., Chang, X.B., Hou, Y.X., He, L., Hegedus, T., Gentzsch, M., Aleksandrov, A., Balch, W.E. and Riordan, J.R. (2007) Domain Interdependence in the Biosynthetic Assembly of CFTR. Journal of Molecular Biology, 365, 981–994.

61. Kai, D. and Lukacs, G.L. (2009) Cooperative assembly and misfolding of CFTR domains in vivo. Mol Biol Cell, 20, 1903–1915.

62. Haggie, P.M., Phuan, P.W., Tan, J.A., Xu, H., Avramescu, R.G., Perdomo, D., Zlock, L., Nielson, D.W., Finkbeiner, W.E., Lukacs, G.L. et al. (2017) Correctors and potentiators rescue function of the truncated W1282X-Cystic Fibrosis Transmembrane Regulator (CFTR) translation product. Journal of Biological Chemistry, 292, 771–785.

63. Ostedgaard, L.S., Randak, C., Rokhlina, T., Karp, P., Vermeer, D., Ashbourne Excoffon, K.J. and Welsh, M.J. (2003) Effects of C-terminal deletions on cystic fibrosis transmembrane conductance regulator function in cystic fibrosis airway epithelia. Proc Natl Acad Sci U S A, 100, 1937–1942.

64. Stratford, F.L., Ramjeesingh, M., Cheung, J.C., Huan, L.J. and Bear, C.E. (2007) The Walker B motif of the second nucleotide-binding domain (NBD2) of CFTR plays a key role in ATPase activity by the NBD1-NBD2 heterodimer. Biochem J, 401, 581–586.

65. Gentzsch, M. and Riordan, J.R. (2001) Localization of sequences within the C-terminal domain of the cystic fibrosis transmembrane conductance regulator which impact maturation and stability. Journal of Biological Chemistry, 276, 1291–1298.

66. Gentzsch, M., Aleksandrov, A., Aleksandrov, L. and Riordan, J.R. (2002) Functional analysis of the C-terminal boundary of the second nucleotide binding domain of the cystic fibrosis transmembrane conductance regulator and structural implications. The Biochemical journal, 366, 541–548.

67. Sharma, N., LaRusch, J., Sosnay, P.R., Gottschalk, L.B., Lopez, A.P., Pellicore, M.J., Evans, T., Davis, E., Atalar, M., Na, C.H. et al. (2016) A sequence upstream of canonical PDZ-binding motif within CFTR COOH-terminus enhances NHERF1 interaction. Am J Physiol Lung Cell Mol Physiol, 311, L1170–L1182.

68. Micoud, J., Chauvet, S., Scheckenbach, K.E.L., Alfaidy, N., Chanson, M. and Benharouga, M. (2015) Involvement of the heterodimeric interface region of the nucleotide binding domain-2 (NBD2) in the CFTR quaternary structure and membrane stability. Biochimica et Biophysica Acta - Molecular Cell Research.

69. Laselva, O., Qureshi, Z., Zeng, Z.W., Petrotchenko, E.V., Ramjeesingh, M., Hamilton, C.M., Huan, L.J., Borchers, C.H., Pomes, R., Young, R. et al. (2021) Identification of binding sites for ivacaftor on the cystic fibrosis transmembrane conductance regulator. iScience, 24, 102542.

70. Fiedorczuk, K. and Chen, J. (2022) Mechanism of CFTR correction by type I folding correctors. Cell, 185, 158–168 e111.

71. Martinovich, K.M., Kicic, A., Stick, S.M., Johnsen, R.D., Fletcher, S. and Wilton, S.D. (2022) Investigating the Implications of CFTR Exon Skipping Using a Cftr Exon 9 Deleted Mouse Model. Front Pharmacol, 13, 868863.

72. Veit, G., Roldan, A., Hancock, M.A., Da Fonte, D.F., Xu, H., Hussein, M., Frenkiel, S., Matouk, E., Velkov, T. and Lukacs, G.L. (2020) Allosteric folding correction of F508del and rare CFTR mutants by elexacaftor-tezacaftor-ivacaftor (Trikafta) combination. JCI Insight, 5.

