## Supplementary material for "Systematic deletion of symmetrical *CFTR* exons reveals new therapeutic targets for exon skipping antisense oligonucleotides": Figure S

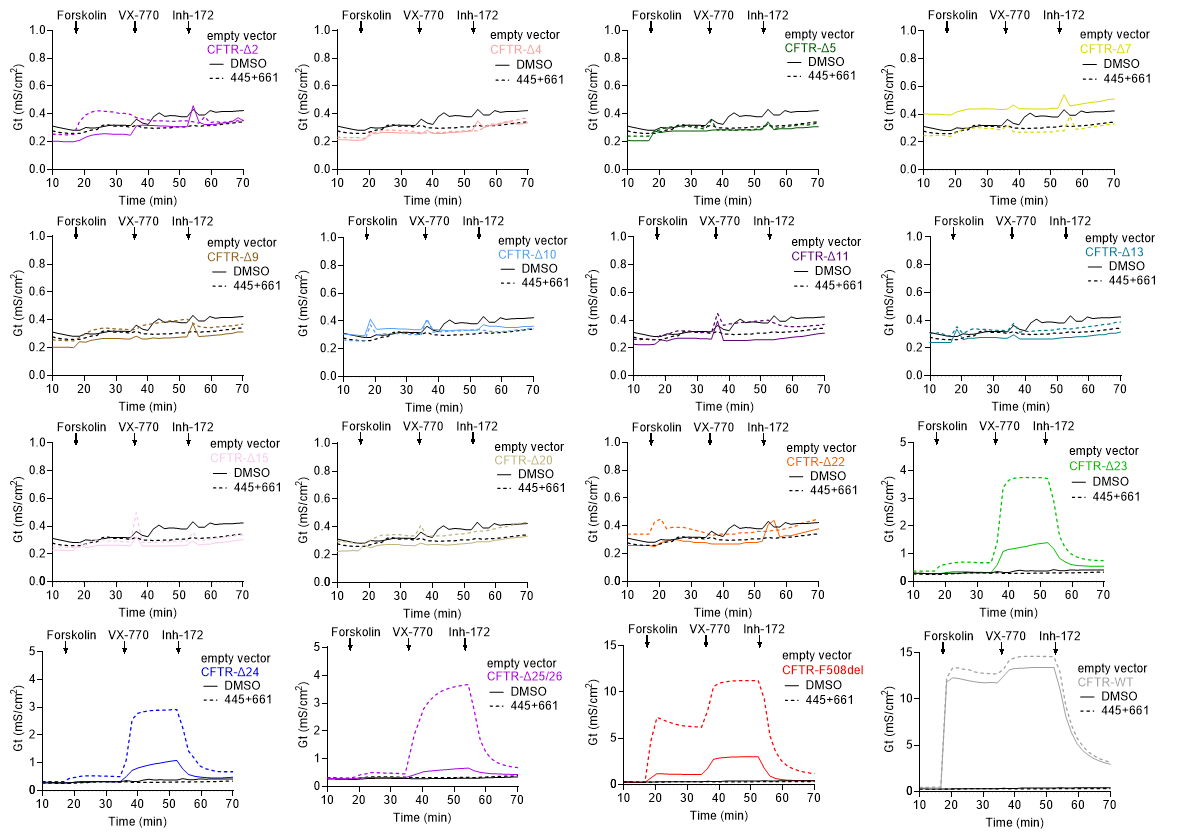

**Figure S1.** Representative current traces of CFTR constructs analyzed in Figure 1C.

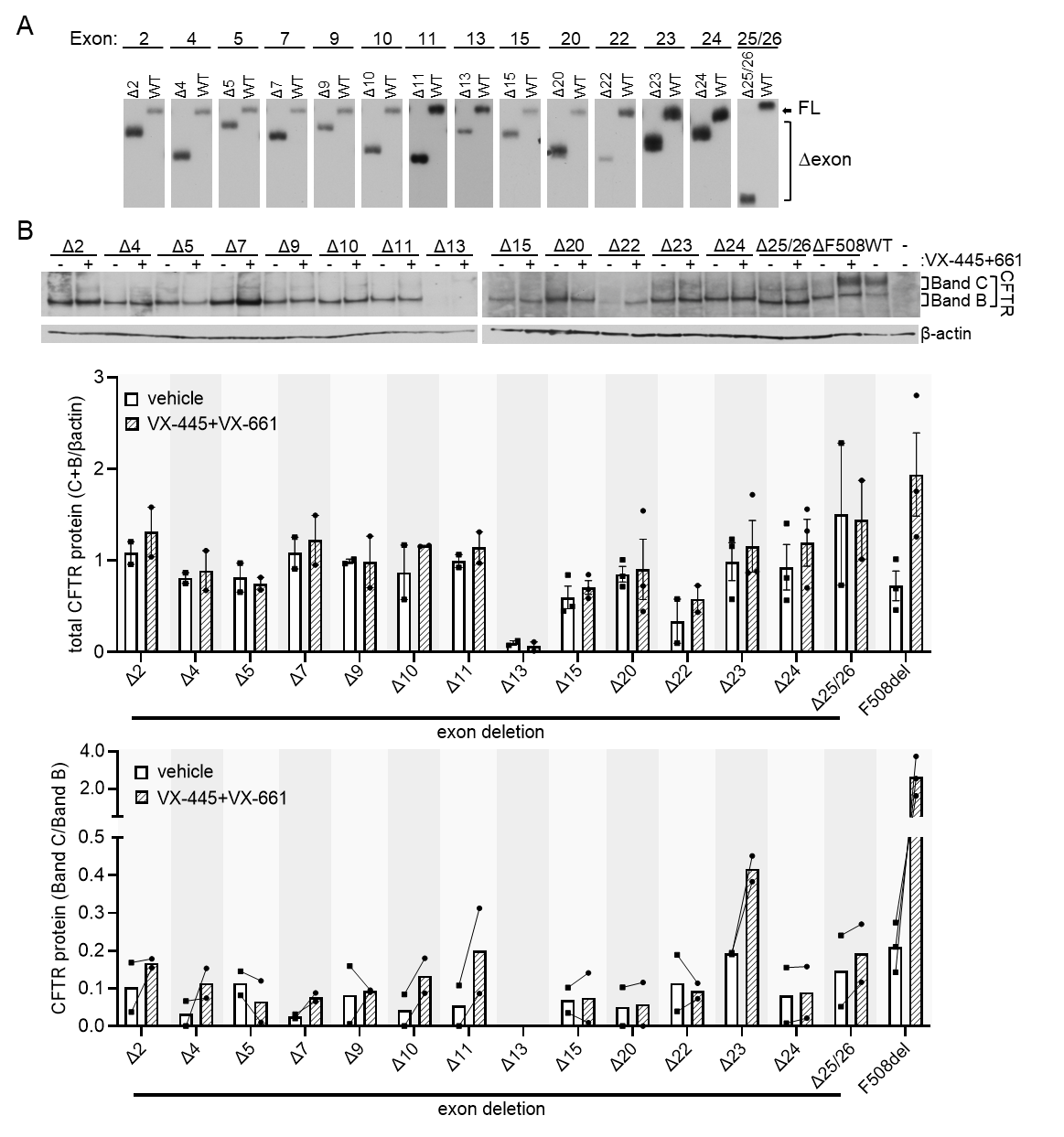

**Figure S2.** Expression of CFTR exon deletion constructs. **(A)** RT-PCR analysis of *CFTR* RNA collected from FRT cells transfected with each CFTR exon deletion construct, compared to CFTR-WT. **(B)** (top) Immunoblot analysis of CFTR protein, bands C and B, isolated from FRT cells stably transfected with empty vector, CFTR-F508del, CFTR-WT, or each CFTR exon deletion construct. Cells were treated with DMSO or VX-445+VX-661. β-actin was used as a loading control. (middle) Total CFTR protein (band C+band B/ β-actin was quantified. (bottom) The ratio of CFTR (band C / band B) isoforms was quantified. Replicates are n=2 and experimental groups are connected by a line.

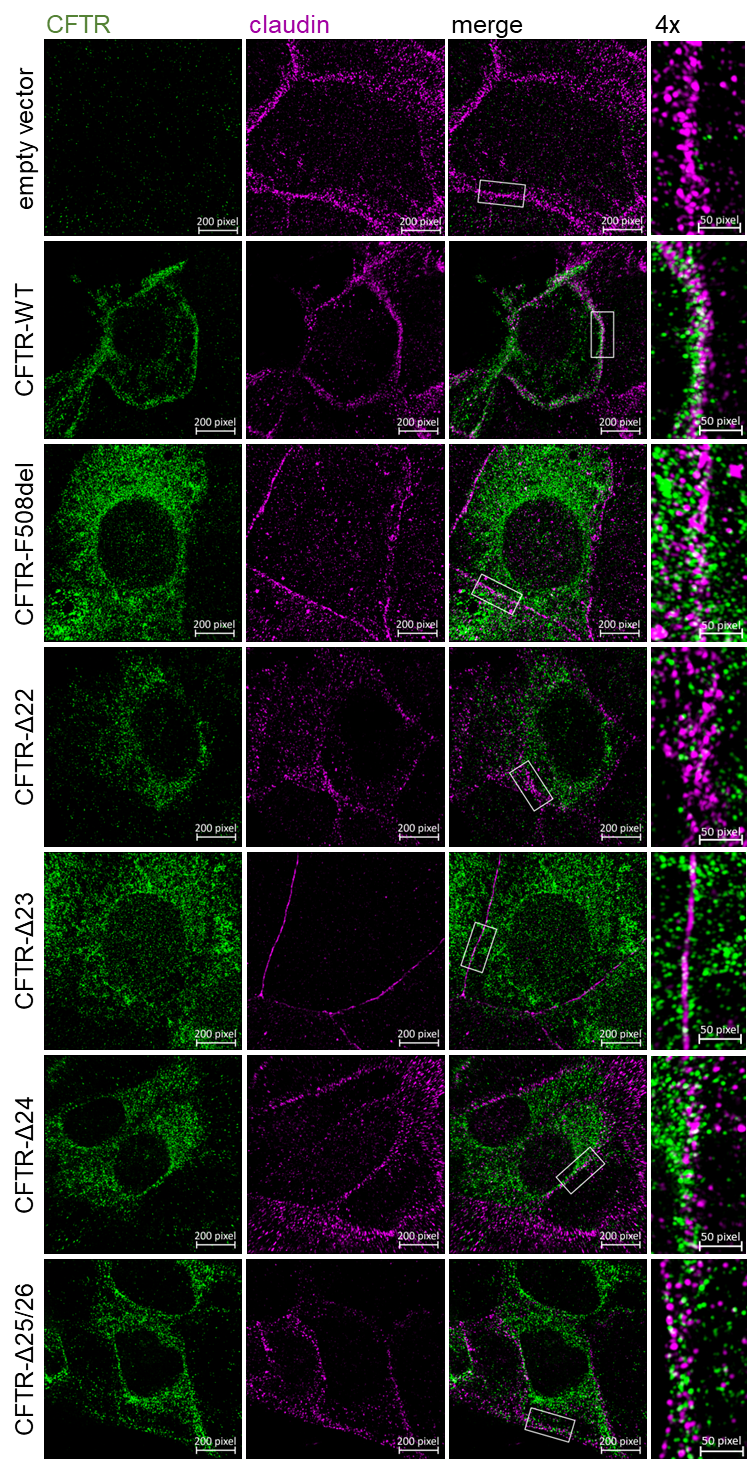

**Figure S3.** Cellular localization of CFTR and the exon skipped isoforms. Immunofluorescent images of FRT cells transfected with empty vector, CFTR-WT, CFTR-F508del, CFTR-Δ22, CFTR-Δ23, CFTR-Δ24, or CFTR-Δ25/26. Cells were incubated with a CFTR-specific monoclonal antibody (green) and a rabbit Claudin polyclonal antibody (magenta). Regions of interest at the cell membrane were enlarged to show co-localization of CFTR at the cell membrane (white).

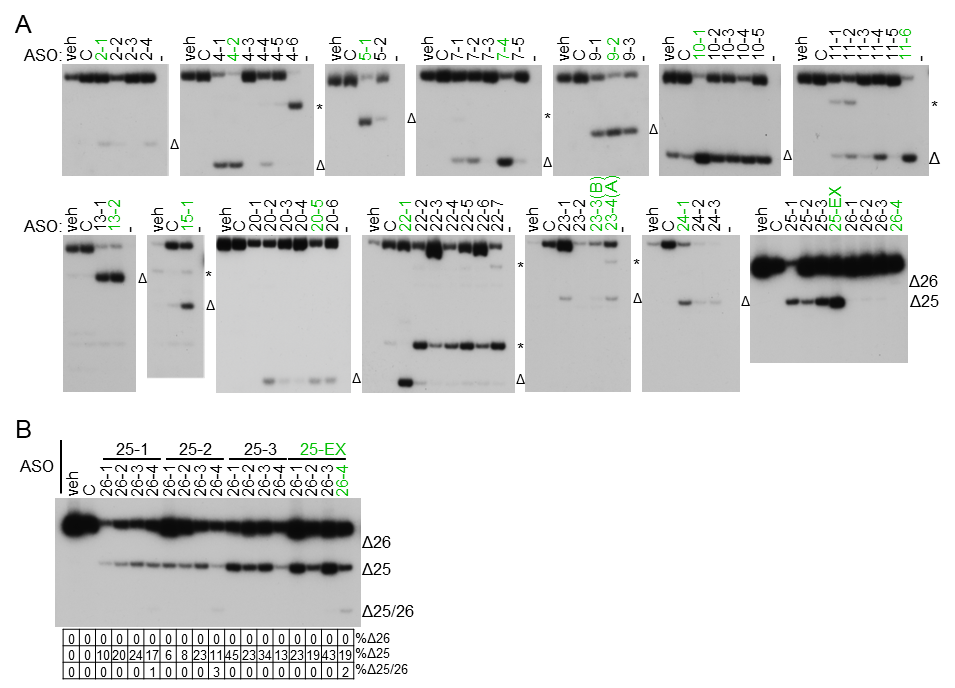

**Figure S4.** Spliced-switching ASOs induce skipping of symmetrical CFTR exons. (**A**) RT-PCR analysis of T84 cells transfected with vehicle, control ASO (ASO-C), or CFTR exon targeting ASOs. Amplicons representing cryptically spliced CFTR is indicated by * and expected exon skipping for each ASO is labelled with Δ on the right side of the gel. Quantification of gels are shown in **Figure 2B**. ASOs that induced the highest level of skipping are highlighted in green. (**B**) RT-PCR analysis of T84 cells transfected with vehicle, control ASO (ASO-C), or CFTR exon 25 and 26 targeting ASOs. Exon skipping is quantified as [Δ/(full-length+Δ)]*100. Primers flanking the exon were used for analysis (**Table S3**).

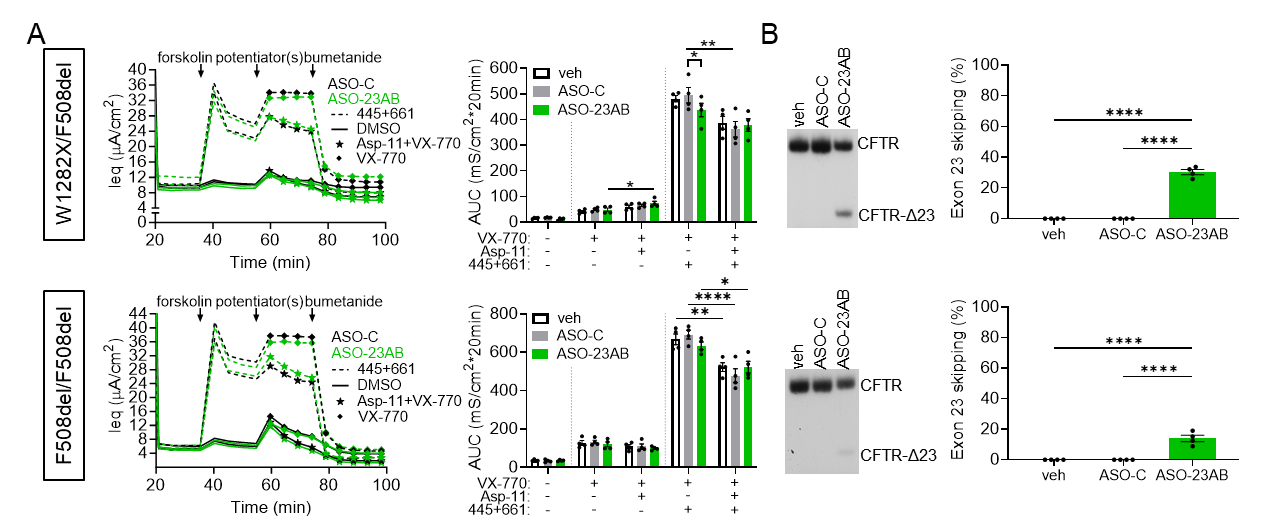

**Figure S5.** ASO treatment induces *CFTR-W1282X* exon skipping but does not increase function above approved HEMTs when expressed with CFTR-F508del (**A**) (left) Equivalent current (Ieq) traces of primary hBE cells from CF donors. The genotype of each donor is indicated on the left. Cells were transfected with ASO-C (80 μM, black), or ASO-23AB (40 μM each, green) and treated with vehicle (DMSO, solid lines) or VX-445+VX-661 (dashed lines) for 24 hours. Forskolin, potentiator [VX-770 (diamond) or VX-770+Asp-11(star)], and bumetanide additions are indicated. (right) Average AUC of the current traces was quantified for the forskolin or forskolin+potentiator(s) test periods for each treatment group. Error bars are ±SEM two-way ANOVA with Tukey’s multiple comparison within groups to assess ASO effects or Šídák's multiple comparison test between groups to assess ASP-11 effects; *P<0.05, **P<0.01.(**B**) (left) RT-PCR analysis of exon 23 splicing in hBE cells from A. (right) Quantification of percent exon 23 skipping for each treated donor. Error bars are ±SEM; one-way ANOVA with Tukey’s multiple comparisons test; *P<0.05, **P<0.01, ****P<0.0001.

| Primers | Sequence (5’-3’) |
| --- | --- |
| 1:HCAI-CFTRdel2R | CTGAAGAACAGCTTGCTCACCAC |
| 2:HCAI-CFTRdel2F | CGAGTGGGACCGCGAGCT |
| 3:HCAI-CFTRdel4R | GCCCAGGTACAGGAAGATG |
| 4:HCAI-CFTRdel4F | ACCCTGAAGCTGAGCAGC |
| 5:HCAI-CFTRdel5R | CTTCTTGTAGATCAGGCTGAACATGGCGATGC |
| 6:HCAI-CFTRdel5F | GGCCTGGCCCTGGCCCAC |
| 7:HCAI-CFTRdel7R | CGGTACTTCATCATCATGC |
| 8:HCAI-CFTRdel7F | GACCGAGCTGAAGCTGAC |
| 9:HCAI-CFTRdel9R | CTGGATCTTGTTGATGGCGC |
| 10:HCAI-CFTRdel9F | GGCTTCGGCGAGCTGTTC |
| 11:HCAI-CFTRdel10R | CTCCTCCCAGAAGGCGGT |
| 12:HCAI-CFTRdel10F | ACCAGCCTGCTGATGGTG |
| 13:HCAI-CFTRdel11R | CTTGCCGGCGCCGGTGCT |
| 14:HCAI-CFTRdel11F | GACATCAGCAAGTTCGCCGAGAAGGACAAC |
| 15:HCAI-CFTRdel13R | CGGGCCAGGCTGATGCGG |
| 16:HCAI-CFTRdel13F | CTGCGTGTGCAAGCTGATGGC |
| 17:HCAI-CFTRdel15R | CTTCAGGTCCTCCTCGTTGATCTCC |
| 18:HCAI-CFTRdel15F | GTGGCCGCCAGCCTGGTG |
| 19:HCAI-CFTRdel20R | CCTCGCTCTCCAGCTGCTTCAGCTGC |
| 20:HCAI-CFTRdel20F | GCGAGGGCGAGGGCCGCG |
| 21:HCAI-CFTRdel22R | CAGGCTGTCCACGTCGATGCTGC |
| 22:HCAI-CFTRdel22F | GTGGGCCTGCTGGGCCGC |
| 23:HCAI-CFTRdel23R | GCGCTGGCCGGGGCTGAT |
| 24:HCAI-CFTRdel23F | AAGGTGTTCATCTTCAGCGGCACCTTC |
| 25:HCAI-CFTRdel24R | CTGGGGGATCACGCCGAAGG |
| 26:HCAI-CFTRdel24F | GTGGGCCTGCGCAGCGTG |
| 27:HCAI-CFTRdel2526R | GTGATCGAGGAGAACAAGGTGC |
| 28:HCAI-CFTRdel2526F | CTCGTCGGCCACCTTCCA |

**Table S1.** Primers flanking *CFTR* symmetrical exons used to generate the *CFTR* exon deletion constructs.

| ASOs | Sequence (5’-3’) | ASOs | Sequence (5’-3’) |
| --- | --- | --- | --- |
| 2-1 | ATCCTTTCCTCAAAATTGGTCTGGT | 13-2 | GGTATTCAAAGAACATACCTTTCAA |
| 2-2 | GTATATGTCTGACAATTCCAGGCGC | 15-1 | ACAATAGAACATTCTTACCTCTGCC |
| 2-3 | CAGATAGATTGTCAGCAGAATCAAC | 20-1 | CAAGATGAGTGAAAATTGGACTCCT |
| 2-4 | GTACATGAACATACCTTTCCAATTT | 20-2 | CGAAGGCACGAAGTGTCCATAGTCC |
| 4-1 | GAGGCTGTACTGCTTTGGTGACTTC | 20-3 | AACAGAGTTTCAAAGTAAGGCTGCC |
| 4-2 | GAAGCTATGATTCTTCCCAGTAAGA | 20-4 | AGTTGGCAGTATGTAAATTCAGAGC |
| 4-3 | GTGTAGGAGCAGTGTCCTCACAATA | 20-5 | TTCTATTCTCATTTGGAACCAGCGC |
| 4-4 | AATGTGATGAAGGCCAAAAATGGCT | 20-6 | GGTAACAGCAATGAAGAAGATGACA |
| 4-5 | GCTATTCTCATCTGCATTCCAATGT | 22-1 | ATGTCAATGAACTTAAAGACTCGGC |
| 4-6 | CCTGTGCAAGGAAGTATTACCTTCT | 22-2 | GGCCAGATGTCATCTTTCTTCACGT |
| 5-1 | CTAGAACACGGCTTGACAGCTTTAA | 22-3 | ATCTTTGACAGTCATTTGGCCCCCT |
| 5-2 | TGGAAAGGAGACTAACAAGTTGTCC | 22-4 | CCACCTTCTGTGTATTTTGCTGTGA |
| 7-1 | ACTGATCTTCCCAGCTCTCTGATCT | 22-5 | TCTCTAATATGGCATTTCCACCTTC |
| 7-2 | ATTTCTGAGGTAATCACAAGTCTTT | 22-6 | CCAGGACTTATTGAGAAGGAAATGT |
| 7-3 | AGTATGCCTTAACAGATTGGATATT | 22-7 | AAGCAGTGTTCAAATCTCACCCTCT |
| 7-4 | ATTTTTTCCATTGCTTCTTCCCAGC | 23-1 | ATCCAGTTCTTCCCAAGAGGCCCAC |
| 7-5 | ATTGGAACAACTTACTGTCTTAAGT | 23-2 | AGCTGATAACAAAGTACTCTTCCCT |
| 9-1 | TCCATCACTACTTCTGTAGTCGTTA | 23-B | AAGTTATTGAATCCCAAGACACACC |
| 9-2 | CTCCTCCCAGAAGGCTGTTACATTC | 23-A | CTAAGTCCTTTTGCTCACCTGTGGT |
| 9-3 | TTAAAAATTCTGACCTCCTCCCAGA | 24-1 | GATCACTCCACTGTTCATAGGGATC |
| 10-1 | GGCTGTCATCACCATTAGAAGTTTT | 24-2 | CTCATCTGCAACTTTCCATATTTCT |
| 10-2 | AATTACTGAAGAAGAGGCTGTCATC | 24-3 | ATTTCAGTTAGCAGCCTTACCTCAT |
| 10-3 | TAATATCTTTCAGGACAGGAGTACC | 25-1 | AGCCCAACCTGCAGTTAGAGAAGAT |
| 10-4 | GATCCAGCAACCGCCAACAACTGTC | 25-2 | TCTTCGCCTTACTGAGAACAGATCT |
| 10-5 | AGAACAAAAGAACTACCTTGCCTGC | 25-3 | AGAACATCTGAAACTCACACTGGAT |
| 11-1 | CTCCCATAATCACCATTAGAAGTGA | 25-EX | CTGGATCCAAATGAGCACTGGGTTA |
| 11-2 | ATTTTACCCTCTGAAGGCTCCAGTT | 26-1 | TACTGTGCAATCAGCAAATGCTTGT |
| 11-3 | ACAGAATGAAATTCTTCCACTGTGC | 26-2 | CCTGTGTTCACAGAGAATTACTGTG |
| 11-4 | GTGCCAGGCATAATCCAGGAAAACT | 26-3 | TCCAGCATTGCTTCTATCCTGTGTT |
| 11-5 | ATGCTTTGATGACGCTTCTGTATCT | 26-4 | GTTATAAAGACTCACCAAAAATTGT |
| 11-6 | TTTTCACATAGTTTCTTACCTCTTC | ASO-C | CCTCTTAGGTCAGTTACAATTTATA |
| 13-1 | TCTAGGTATCCAAAAGGAGAGTCTA |  |  |

**Table S2.** Antisense oligonucleotide sequences used in this study.

| Primers | Sequence (5’-3’) | Primers | Sequence (5’-3’) |
| --- | --- | --- | --- |
| 1:HCAI-CFTR-ex1F | ATGCAGCGCAGCCCCCTC | 25:hCFTR-ex10F | CTAATGGTGATGACAGCCTC |
| 2:HCAI-CFTR-ex6R | CGGTACTTCATCATCATGC | 26:hCFTR-ex12R | GCTCGTTGACCTCCACTCAG |
| 3:HCAI-CFTR-ex5F | ACCCTGAAGCTGAGCAGC | 27:hCFTR-ex12F | CACTGAGTGGAGGTCAACGA |
| 4:HCAI-CFTR-ex9R | CTCCTCCCAGAAGGCGGT | 28:hCFTR-ex14R | TCCAGGAGACAGGAGCATCT |
| 5:HCAI-CFTR-ex8F | GACCGAGCTGAAGCTGAC | 29:hCFTR-ex14F | AGGAGGCAGTCTGTCCTGAA |
| 6:HCAI-CFTR-ex11R | CTCCTCCAGCTGGCAGGC | 30:hCFTR-ex16R | CAGCACAACCAAAGAAGCAG |
| 7:HCAI-CFTR-ex9F | GACTTCCTGCAGAAGCAG | 31:hCFTR-ex19F | CAGTGCCAGTGATAGTG |
| 8:HCAI-CFTR-ex14RB | CATCAGCTTGCACACGCA | 32:hCFTR-ex21R | CTCTTCCTTCTCCTTCTC |
| 9:HCAI-CFTR-ex14FE | GCAAAGTGAGCCTGGCCC | 33:hCFTR-ex21F | GGGCTGTAAACTCCAGCATA |
| 10:HCAI-CFTR-ex19R | CCTCGCTCTCCAGCTGCTTCAGCTGC | 34:hCFTR-ex23R | GACACACCATCGATCTGG |
| 11:HCAI-CFTR-ex21F | GCGAGGGCGAGGGCCGCG | 35:hCFTR-ex22F | CCAAACCATACAAGAAT |
| 12:HCAI-CFTR-ex25R | ACGGGGTCCAGGTGGGCG | 36:hCFTR-ex24R | GATCACTCCACTGTTCAT |
| 13:hCFTR-ex1F | GCAGGCACCCAGAGTAGTAG | 37:hCFTR-ex23F | GGGAAGAGTACTTTGTTATC |
| 14:hCFTR-ex3R | GCCAGCTCTCTATCCCATTC | 38:hCFTR-ex25R | GTTCTATCACAGATCTGAG |
| 15:hCFTR-ex3F | GGGATAGAGAGCTGGCTTCA | 39:hCFTR-ex24F | CTTGGATCCCTATGAACA |
| 16:hCFTR-ex5R | TTGTTCAGGTTGTTGGAAAGG | 40:hCFTR-ex27R | CTAAAGCCTTGTATCTTG |
| 17:hCFTR-ex4F | CAAGGAGGAACGCTCTATCG | 41:hβ-actinFor | AAAGACCTGTACGCCAACAC |
| 18:hCFTR-ex6R | AGGCAGACGCCTGTAACAAC | 42:hβ-actinRev | GTCATACTCCTGCTTGCTGAT |
| 19:hCFTR-ex6F | ATGGGGCTAATCTGGGAGTT | 43:qhCFTR-ex11WTF | TGGCACCATTAAAGAAAATATCATCTT |
| 20:hCFTR-ex8R | CTGCCTTCCGAGTCAGTTTC | 44:qhCFTR-ex12WTR | CTCAGTGTGATTCCACCTTCTC |
| 21:hCFTR-ex8F | TCACCACCATCTCATTCTGC | 45:qhCFTR-ex11ΔFF | GGCACCATTAAAGAAAATATCATTGG |
| 22:hCFTR-ex10R | AAGAAGAGGCTGTCATCACCA | 46:qhCFTR-ex12ΔFR | CTCAGTGTGATTCCACCTTCT |
| 23:hCFTR-ex9F | TAACAGCCTTCTGGGAGGAG | 47:qhHPRT1For | GCGATGTCAATAGGACTCCAG |
| 24:hCFTR-ex11R | GGCATGCTTTGATGACGCTT | 48:qhHPRT1Rev | TTGTTGTAGGATATGCCCTTGA |

| Probes | Sequence (5’-3’) |
| --- | --- |
| hCFTR-DF508 | 56-FAM/ACAGAAGCG/ZEN/TCATCAAAGCATGCC/3IABkFQ |
| hCFTR-F508 | 5.6-FAM/ACAGAAGCG/ZEN/TCATCAAAGCATGCC/3IABkFQ |
| hHPRT1 | 5HEX/AGCCTAAGA/ZEN/TGAGAGTTCAAGTTGAGTTTGG/3IABkFQ |

**Table S3.** Primers and probes used for PCR and qPCR analysis.
